# A zinc finger transcriptional repressor ZOS5-09 regulates rice grain size, starch and protein biosynthesis

**DOI:** 10.1101/2024.05.26.595847

**Authors:** Priya Jaiswal, Falah Qasim, Arunima Mahto, Ankur Vichitra, Akhilesh K. Tyagi, Pinky Agarwal

**Affiliations:** National Institute of Plant Genome Research. New Delhi, India; Interdisciplinary Centre for Plant Genomics and Department of Plant Molecular Biology, University of Delhi South Campus, New Delhi, India

**Author notes:** Address correspondence to /.

**Keywords:** C_2_H_2_ zinc finger transcription factor, grain, rice, seed storage protein, starch, transcriptional repressor

## Abstract

**Highlight:** A rice seed-preferential transcriptional repressor, ZOS5-09 controls grain size by affecting cell proliferation and expansion; and regulates starch and protein biosynthesis. It acts as a repressor by deacetylation.

Grain size is the one of the key determinants of grain yield. Transcription factors form an important class of regulatory proteins that play crucial roles in regulating developmental responses. Our study focuses on a novel seed-preferential C_2_H_2_ zinc finger transcription factor, ZOS5-09 (LOC_Os05g38600) that plays an important role in regulating grain related traits in rice. Rice plants with *ZOS5-09* promoter::*GUS* construct showed its high expression in rice endosperm. *In planta* reporter effector assays and localization studies showed that ZOS5-09 is a nuclear localized transcriptional repressor. It has a C-terminal NoRS (nucleolar retention signal). Ectopic and seed-preferential overexpression of *ZOS5-09* resulted in lethality. Seed-preferential overexpression without NoRS was detrimental to grain filling. Rice plants with knock-down or CRISPR based knock-out of *ZOS5-09* displayed reduced grain length and weight but increased grain width. Plant height, second leaf width, and panicle number were reduced and second leaf length were increased. The change in grain size was due to lower cell proliferation and increased cell size in the transverse direction due to down regulation of cell cycle related genes and increased expression of expansins. Decreased expression of *ZOS5-09* also resulted in reduced total starch and protein content and higher endosperm chalkiness, thus, negatively affecting grain quality. ZOS5-09 directly regulated a seed storage protein encoding gene, *GLU6* by binding to the zinc finger binding site in its promoter. It acted as a repressor by promoting deacetylation upon interaction with a histone deacetylase. In summary, our results indicate that optimum expression of *ZOS5-09* is essential for proper rice grain development. Our study highlights the role of a transcriptional repressor in regulating rice grain traits and improves our understanding of the transcriptional regulatory networks affecting grain size.

## Introduction

Plant growth and development are regulated by an intricate network of transcription factors (TFs) which act in a coordinated manner to attain a fine regulatory control. Diverse TF families like NAC, WRKY, Zn finger, bHLH, bZIP, MADS, and AP2-ERF are known to regulate various plant developmental processes. Zinc finger (ZF) TF is one of the largest families, which includes nine subfamilies based on Cys and His motifs (Schumann *et al*., 2007). C_2_H_2_ ZF protein is the largest TF subfamily that acts as a key regulator of transcriptional processes. The first identified C_2_H_2_ ZF protein was *Petunia* EPF1. It binds to the promoter of *5-enolpyruvylshikimate-3-phosphate synthase* and regulates the expression of floral homeotic genes (Takatsuji *et al*., 1992). Most of the C_2_H_2_ ZF proteins contain a QALGGH motif in the ZF region and are known as Q-type. Those lacking QALGGH motif are called C-type. C_2_H_2_ ZF proteins are also known as TFIIIA-type proteins and are well-characterized in various plant species. They play a wide variety of functions in regulating plant developmental processes. In addition, they also play important roles in biotic and abiotic stresses (Jiang and Pan, 2012, Han *et al*., 2020). Till now, 189 C_2_H_2_ ZF proteins have been reported in *Oryza sativa* (Agarwal *et al*., 2007), 122 in *Triticum durum* (Faraji *et al*., 2018), 321 in *Glycine max* (Yuan *et al*., 2018a), 301 in *Brassica rapa* (Alam *et al*., 2019), 104 in *Solanum lycopersicum* (Hu *et al*., 2019), 218 in *Medicago truncatula* (Jiao *et al*., 2020), 129 in *Cucumis sativa* (Yin *et al*., 2020), 79 in *Solanum tuberosum* (Liu *et al*., 2020), 98 in *Vitis vinifera* (Arrey-Salas *et al*., 2021) and 158 in *Arabidopsis thaliana* (Li *et al*., 2022).

Plant C_2_H_2_ ZF proteins often act as transcriptional repressors and suppress target gene expression. Most of the plant transcriptional repressors harbor ethylene-responsive element binding factor-associated amphiphilic repression (EAR) motif for their repression activity (Kagale and Rozwadowski, 2011). The repressive activity of the EAR motif is decided by the consensus sequence patterns LxLxL or DLNxxP (Singh *et al*., 2019). Such proteins play crucial roles in regulating plant development, hormone signaling and response to external stimuli (Wang *et al*., 2020c). Some such C_2_H_2_ ZF proteins like SUPERMAN and GLABROUS INFLORESCENCE STEMS3 (GIS3) regulate flower and fruit development in *Arabidopsis thaliana* respectively (Sun *et al*., 2015, Rodas *et al*., 2021). In rice, OsZFP207 regulates gibberellin biosynthesis (Duan *et al*., 2021). Drought-responsive ZF protein (OsDRZ1) regulates growth and drought tolerance (Yuan *et al*., 2018b). EAR motif containing proteins interact with co-repressors like TOPLESS (TPL), TOPLESS related proteins (TPRs) and histone modifiers (histone deacetylases or histone demethylase) to repress gene expression of downstream genes (Chang *et al*., 2019, Wang *et al*., 2020a, Fang *et al*., 2021). SlERF.F12 containing an EAR motif, interacts with TPL2 and HDA1 to regulate fruit ripening in tomato (Deng *et al*., 2022). BRI1-EMS-SUPPRESSOR 1 (BES1) is another EAR containing transcriptional regulator that interacts with TPL and HDA19 to regulate BR signaling in *Arabidopsis* (Kim *et al*., 2019). Hence, transcriptional repression plays a crucial role in fine tuning of gene expression and regulating developmental processes.

Seed development is a complex, continuous transcriptional programme wherein the embryo and the endosperm development occur in a sequential manner (Counce and Moldenhauer, 2019, Mahto *et al*., 2023). TFs control gene expression in a time-specific manner to regulate this process. NON STOP GLUMES1 encodes for a C_2_H_2_ ZF TF and is known to regulate spikelet development in rice (Zhuang *et al*., 2020). Tipped1 is a C_2_H_2_ ZF transcriptional repressor that regulates awn development in *Triticum durum* (Huang *et al*., 2020a). AtZAT4 has been reported to regulate seed development in *Arabidopsis* (Puentes-Romero *et al*., 2022).

In this study, we delineate the molecular and functional roles of a seed-preferential C_2_H_2_ ZF TF, ZOS5-09, which acts as a transcriptional repressor and plays an important role in reproductive development in rice. Rice *pZOS5-09::GUS* transgenic plants show an endosperm-specific expression directed by the promoter. Phenotyping of knock-down rice plants and CRISPR knock-out mutants of *ZOS5-09* show decreased grain length, grain weight and increased grain width. The starch, amylose and total protein content also decrease in these grains. ZOS5-09 directly binds to the zinc finger binding site on the promoter of *GLU6* to down regulate it. The gene is highly detrimental to plant growth when ectopically or seed-preferentially overexpressed. Seed preferential overexpression without the nucleolar retention signal allows the plant to reach till the panicle stage, but prevents grain formation. ZOS5-09 represses downstream genes by deacetylation. The study concludes that *ZOS5-09* is an important regulator of rice grain development.

## Results

### Expression analysis of *ZOS5-09*

*ZOS5-09* was found to highly express in rice seed development stages by microarray analyses (Sharma et al, 2012 and Singh et al., 2019). Quantitative real time PCR (qRT-PCR) based expression profiling of *ZOS5-09* validated the same (Supplementary Fig. S1A). The levels of expression gradually increased from S1 to S4, and reached the highest at S5 (∼575 folds). *ZOS5-09* exhibited low levels of expression in P1 panicle stage, callus, and vegetative tissues like root, node, internode, leaf sheath and second leaf. A 1618 bp upstream promoter region of *ZOS5-09* revealed the presence of six elements (CMSRE1IBSPOA, DPBFCOREDCDC3, MYBCORE, PYRIMIDINEBOXOSRAMY1A, RAV1AAT and Skn-1_motif), which could be responsible for its seed-preferential expression. Rice transgenic plants harboring the *pZOS5-09::GUS* construct were used to study the functionality of the promoter. The presence of blue color indicated mild expression in calli, but no expression in leaf, root, pre-pollination floret and S1-S2 stages of seed development (Fig. 1A-1-6). S1 stage is 0-2 days after pollination (DAP) while 3-4 days pollinated seeds are considered as S2 stage. *GUS* expression was observed in S3, S4, and S5 stages of developing seed, specifically in the endosperm (Fig. 1A-7, 8). S3 stage is 5-10 DAP, S4 stage is 11-20 DAP and S5 is 21-30 DAP. In S5 stage, expression was extremely high in the outer layers of endosperm (Fig. 1A-9). The result was corroborated through *GFP* expression analysis of rice plants transformed with *pZOS5-09::GFP.* Mild *GFP* expression was observed in callus and S1 and S2 seed stages (Supplementary Fig. S1B-1-5, 13). The *GFP* expression in the endosperm was clearly visible at the S3 stage (Supplementary Fig. S1B-6-9), but was less distinguishable from the WT at S4 due to background fluorescence (Supplementary Fig. S1B-10, 12). Interestingly, mild expression was also observed in the radicle region of embryo at S3 and S4 stages in the plants containing *GFP* reporter gene (Supplementary Fig. S1B-9, 11). It may be due to the expression of this gene specifically, in the radicle region cells, present in the inner layers of the embryo. The transgenic rice seeds containing the *GUS* reporter gene, however, did not show GUS expression in the embryo region of S3 and S4 stages. The results also reflect on the relative advantages of GUS and GFP reporter genes for studying promoter directed expression.

**Fig. 1.**
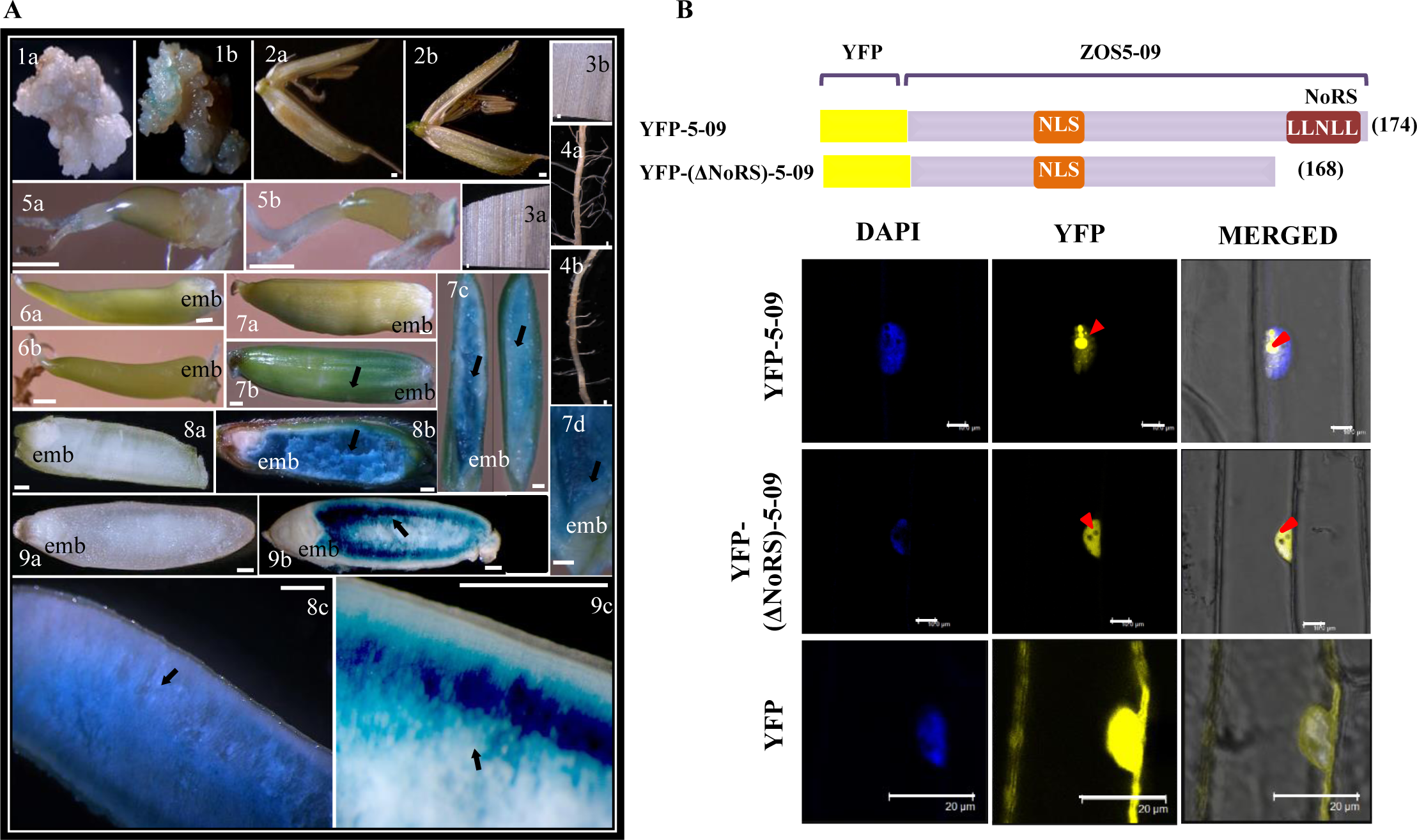
Expression analysis and sub-cellular localization of ZOS5-09. (A) GUS expression from *ZOS5-09* promoter in *pZOS5-09::GUS* rice transgenic plants (all ‘b’ figures) as compared to wild type (all ‘a’ figures) in (1a, b) callus; (2a, b) pre-pollination floret; (3a, b) leaf; (4a, b) root; (5a, b) seed stage S1; (6a, b) seed stage S2; (7a, b) seed stage S3; (7c) transgenic S3 endosperm and embryo; (7d) transgenic S3 embryo; (8a, b) seed stage S4; (8c) transgenic S4 endosperm; (9a, b) seed stage S5; (9c) transgenic S5 endosperm. Scale bar is 0.5 mm. embryo (emb) position has been marked. Black arrows indicate GUS expression. (B) Schematic representation of the full length ZOS5-09 protein fused to YFP. NLS and NoRS represent nuclear localization and nucleolar retention signals, respectively. YFP-(ΔNoRS)-5-09 represents protein with NoRS removed. The numbers in bracket indicate the protein length. Confocal microscopy images show the localization of the YFP fusion constructs in onion epidermal cells. Left panel represents 4′,6-diamidino-2-phenylindole (DAPI) fluorescence staining of the nucleus at 358 nm. Middle panel represents YFP fluorescence at 514 nm. Right panel represents both images merged with bright field images to show sub-cellular localization. Red arrow indicates nucleolus.

#### Nuclear localized ZOS5-09 functions as a transcriptional repressor

In terms of subcellular localization, in both onion peel and *Nicotiana benthamiana* leaf cells, full-length ZOS5-09 (YFP-5-09) was localized to the nucleus with a strong signal in the nucleolus (Fig. 1B, Supplementary Fig. S2). On deletion of nucleolar retention signal (NoRS), YFP-(ΔNoRS)-5-09 was localized to the nucleus, and excluded from the nucleolus (Fig. 1B). This confirmed that ZOS5-09 is a nuclear-localized protein and the C-terminal leucine rich region (LRR) acted as the NoRS. ZOS5-09 has an EAR (an ethylene-responsive element binding factor-associated amphiphilic repression) motif towards the C-terminal (Supplementary Fig. S3A). This is a DLN type of EAR motif ((DLNSPP). On checking for transactivation in yeast, cells transformed with ZOS5-09_pGBKT7 were unable to grow on TDO media and there was absence of blue color on SD/-Trp media supplemented with X-β-gal (similar to negative control), which indicated that ZOS5-09 is not a transcriptional activator (Supplementary Fig. S3B). The DLN motif of ZOS5-09 imparted a transrepression ability in yeast (Singh *et al*., 2019). To confirm its repression property, *in planta* reporter effector assay was carried out in *Nicotiana benthamiana* leaf cells (Supplementary Fig. S3C). For this, *GUS* under the control of *pCys-Prt*, acted as the reporter. It is known that MADS29 binds to this promoter and activates its expression (Yin and Xue, 2012). Cells transformed with only the reporter construct, *pCys-Prt*::*GUS* showed minimal *GUS* expression while cells co-transformed with both the effector and reporter constructs, showed a 415.55% increase in *GUS* expression. When the effector construct, *pCaMV35S*::*MADS29*:*DLN* was co-transformed with the reporter construct, *GUS* expression was lowered by 92.49% and 61.29% with respect to *pCaMV35S*::*MADS29* and *pCys-Prt,* respectively. This showed that DLNSPP motif of ZOS5-09 could repress the activation property of MADS29 on *pCys-Prt* promoter and *in planta* acted as a transcriptional repression motif.

#### Ectopic or seed-preferential overexpression of *ZOS5-09* leads to developmental arrest of the plant

To understand the biological functions of *ZOS5-09* in the growth and development of rice plants, transgenic rice plants ectopically expressing *ZOS5-09* under a *UBIQUITIN* promoter (5-09_OE) were generated. The overexpression of *ZOS5-09* was highly detrimental to plant growth. 5-09_OE calli showed highly decreased plant regeneration ability and mostly turned brown while the control calli turned green on regeneration media (Supplementary Fig. S4A). The presence of the overexpression construct of *ZOS5-09* was confirmed by observance of high levels of reporter gene expression in the selected calli as well as the few plants, obtained on regeneration media (Supplementary Fig. S4B, C, F). qRT-PCR analysis showed ∼7327 to 14503 folds increased expression of *ZOS5-09* in 5-09_OE plants as compared to wild type. 5-09_OE plants exhibited leaf curling and highly retarded growth, turned yellow-brown, senesced and eventually did not survive (Supplementary Fig. S4D-F). Since the transgenic plants had such high levels of *ZOS5-09* expression, the observed phenotype could be attributed to the overexpression of the gene.

Seed preferential overexpression (SOE) of ZOS5-09 (5-09_SOE_1) under *GluD-1* promoter also showed highly decreased plant regeneration ability (Supplementary Fig. S5D). SOE plants of ZOS5-09 without the NoRS sequence (5-09_SOE_2) showed normal plant development but no grain filling (Supplementary Fig. S5E). 5-09_SOE_2 plants showed reduced plant height, increased flag leaf length, and flag leaf width as compared with wild type but other vegetative characters such as number of tillers per plant, stem thickness, second leaf length, second leaf width, panicle morphology, number of panicles per plant and panicle length, were unaltered (Supplementary Fig. S5F-P). Hence, ectopic or seed-preferential overexpression of *ZOS5-09*, leads to lethality. Seed-preferential overexpression of the truncated form with NoRS deleted, prevents grain filling. This implies that excessive levels of *ZOS5*-09 are detrimental to rice plant.

#### Generation of *ZOS5-09* mutants using CRISPR-Cas9 technology in rice

On the basis of the protein structure of ZOS5-09, two sgRNAs, one targeting the region encoding the first ZF domain (ZF) and another targeting the NoRS (SS) coding region were designed (Supplementary Fig. S6A, B). Four independent mutant lines for ZF target knock-out (ZF_5-09) and six independent mutant lines for SS target knock-out (SS_5-09) were generated. Indels were detected in T_0_ transformed lines (shown by chromatogram in Supplementary Fig. S6C, D) and their mutation type was established (Supplementary Fig. S7A). Grain filling in T_0_ transformed lines were highly affected (Supplementary Fig. S7A). Subsequent generations also showed decreased germination efficiency (Supplementary Fig. S7A, S8). Due to extremely low rate of grain filling and germination rate, only ZF_4 and SS_1 lines were proceeded for analysis in T_2_ generation. Sequence analysis of respective target regions (Supplementary Fig. S6E, S7A) showed A insertion in ZF_4 and AA insertion in SS_1 plants. Indels created in both ZF_4 and SS_1 lines led to frameshift mutations resulting in truncated proteins (Supplementary Fig. S6F). The reading frames were changed after seven and 164 amino acids in ZF_4 and SS_1, respectively. To ensure that the resulting phenotype was due to indels created in the coding sequence of *ZOS5-09*, off-target gene validation was done in knock-out plants of *ZOS5-09* and no editing was observed (Supplementary Fig. S7B). The phenotype of these plants has been discussed in the next section.

#### *ZOS5-09* affects grain size through regulation of cell cycle

In addition to knock-out plants, we also generated knock-down (KD) plants for *ZOS5-09* using the RNAi technology. The T_1_ grains on 5-09_KO and 5-09_KD plants showed a decrease in average per grain length and per grain weight but an increase in average per grain width as compared to WT grains (Supplementary Fig. S9, S10). Since the expression levels of *ZOS5-09* were highest in the S5 stage of seed development (Supplementary Fig. S1A), the relative expression levels of *ZOS5-09* were checked at this stage, in the T_2_ generation. *ZOS5-09* was downregulated by ∼3.6 to 7.9 folds in 5-09_KD plants (Supplementary Fig. S11C).

Subsequently, 5-09_KD and 5-09_KO plants showed a decrease in plant height, second leaf width and number of panicles per plant while the second leaf length was increased as compared with WT. Panicle architecture was unaltered (Supplementary Fig. S11A-B, D-G). Further, seed measurements revealed that 5-09_KD and 5-09_KO per grain length was decreased by 0.39-2.34%, per grain width was increased by 0.97-7.28% and per grain weight was decreased by ∼5.5% (Fig. 2A-E). The number of filled grains per plant was decreased in both 5-09_KD and 5-09_KO plants. The number of unfilled grains per plant was significantly increased in 5-09_KD but unaffected in 5-09_KO plants. The overall grain yield was significantly decreased in 5-09_KD by ∼35% and by ∼11% in 5-09_KO plants (Fig. 2F-H). Here, grain yield per plant is referred to the total weight of filled grains per plant. The probable reason for this difference between KD and KO plants maybe the lowered expression of the segmental duplicate of *ZOS5*-09 (Supplementary Fig. S12) only in 5-09_KD plants, which is the single gene sharing more than 50% homology with *ZOS5-09*. Since these are 5-09_KO and 5-09_KD plants, the results suggested that *ZOS5-09* functions as a negative regulator of grain width and a positive regulator of grain length and weight in rice.

**Fig. 2.**
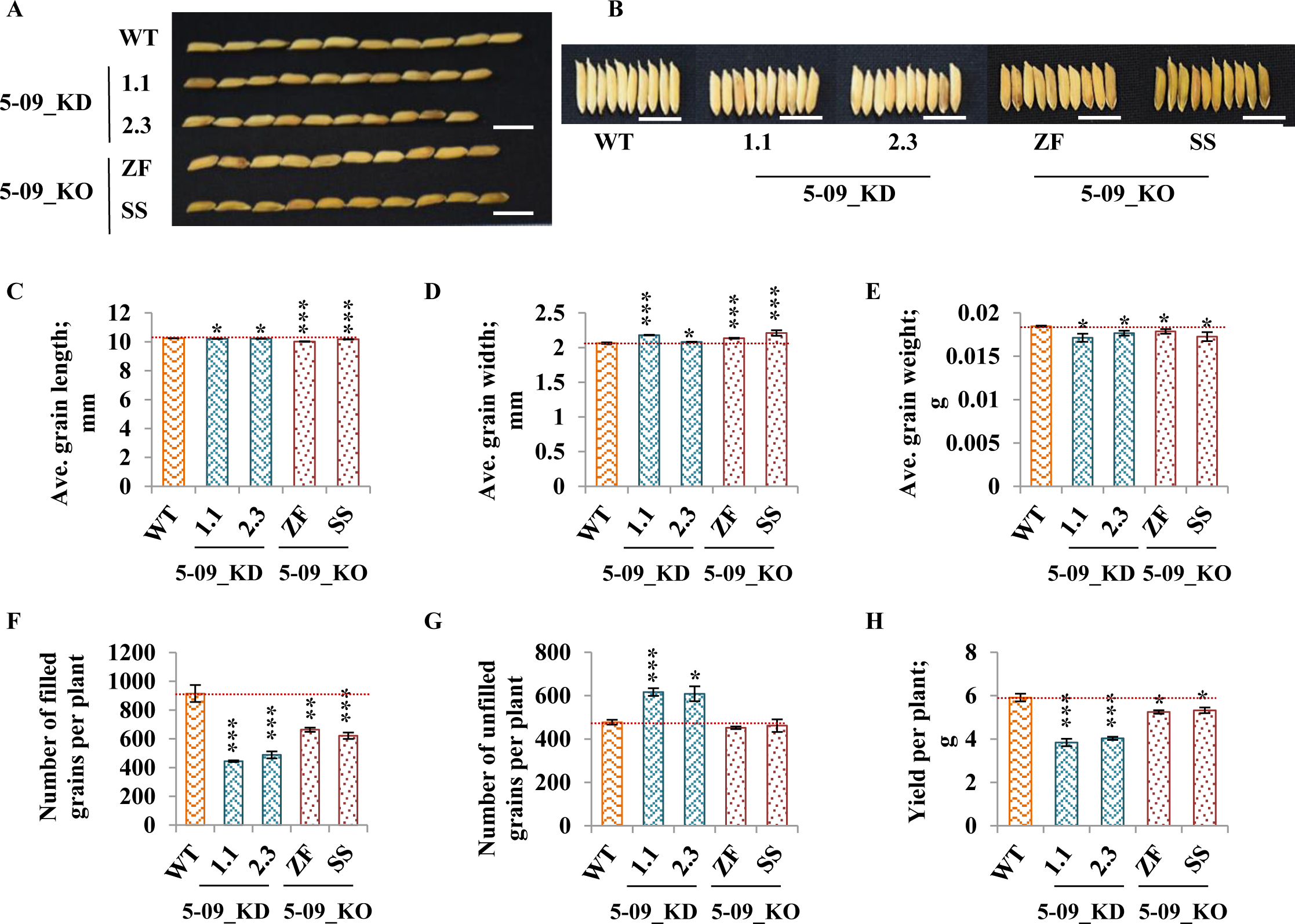
Phenotyping of *ZOS5-09* knock-down (KD) and knock-out (KO) grains. Image of mature rice grains arranged in (A) lengthwise and (B) widthwise directions to show change in grain length and grain width in 5-09_KD T_4_ and 5-09_KO T_3_ grains. (C-E) Bar graphs show (C) average per grain length, (D) average per grain width and (E) average per grain weight (n=50*3 for C-E). (F-H) Bar graphs show (F) change in number of filled grains per plant, (G) number of unfilled grains per plant and (H) yield per plant, respectively, in 5-09_KD/ 5-09_KO plants as compared to WT (n=3 for F-H). Transgenic lines analyzed for ZF and SS are ZF_4 and SS_1 respectively. Statistical significance for 5-09_KD and 5-09_KO have been calculated with respect to WT. Asterisks denote significant difference as determined by Student’s t-test (*,**,*** = p value ≤ 0.05, ≤ 0.01, ≤ 0.005 respectively). Error bars = ± SE.

The lemma cells have a direct forbearance on grain size (Ren *et al*., 2018). The central lemma cells of 5-09_KD and 5-09_KO grains appeared slightly larger than WT (Fig. 3A). The lemma cell number in the longitudinal direction was decreased by ∼25% and the cell length was increased by 18-35% (Fig. 3B-D, Supplementary Fig. S13A). In the transverse direction, representing grain width, the cell number was also decreased by ∼23% but cell width was increased by 28-55% (Fig. 3E-G, Supplementary Fig. S13B). This indicated more cell expansion in the transverse direction as compared with cell expansion in the longitudinal direction, resulting in wider cells (Fig. 3A, Supplementary Fig. S13C). This was confirmed by the decreased ratio of average length between lemma cells in longitudinal over the transverse direction (Supplementary Fig. S13C). In addition, just like lemma cells, the endosperm cell length was also increased in 5-09_KD and 5-09_KO grains by 5.7-19.2% (Fig. 3F, Supplementary Fig. S14) and the number of endosperm cells were significantly decreased in 5-09_KD and 5-09_KO grains as compared to wild type (Fig. 3G, Supplementary Fig. S14). Further, the cell cycle related genes *CDKA1, CYCA, CYCB, CDC20* and *MCM3* were down regulated while expansins *EXPA1*, *EXPA2, EXPA3* and *EXPA4* were up regulated in 5-09_KD and 5-09_KO grains (Fig. 4A, B). This corroborates with reduced cell proliferation and increased cell expansion in 5-09_KD and 5-09_KO grains. Hence, ZOS5-09 regulates both cell proliferation and cell expansion in the rice grain, which affects its size.

**Fig. 3.**
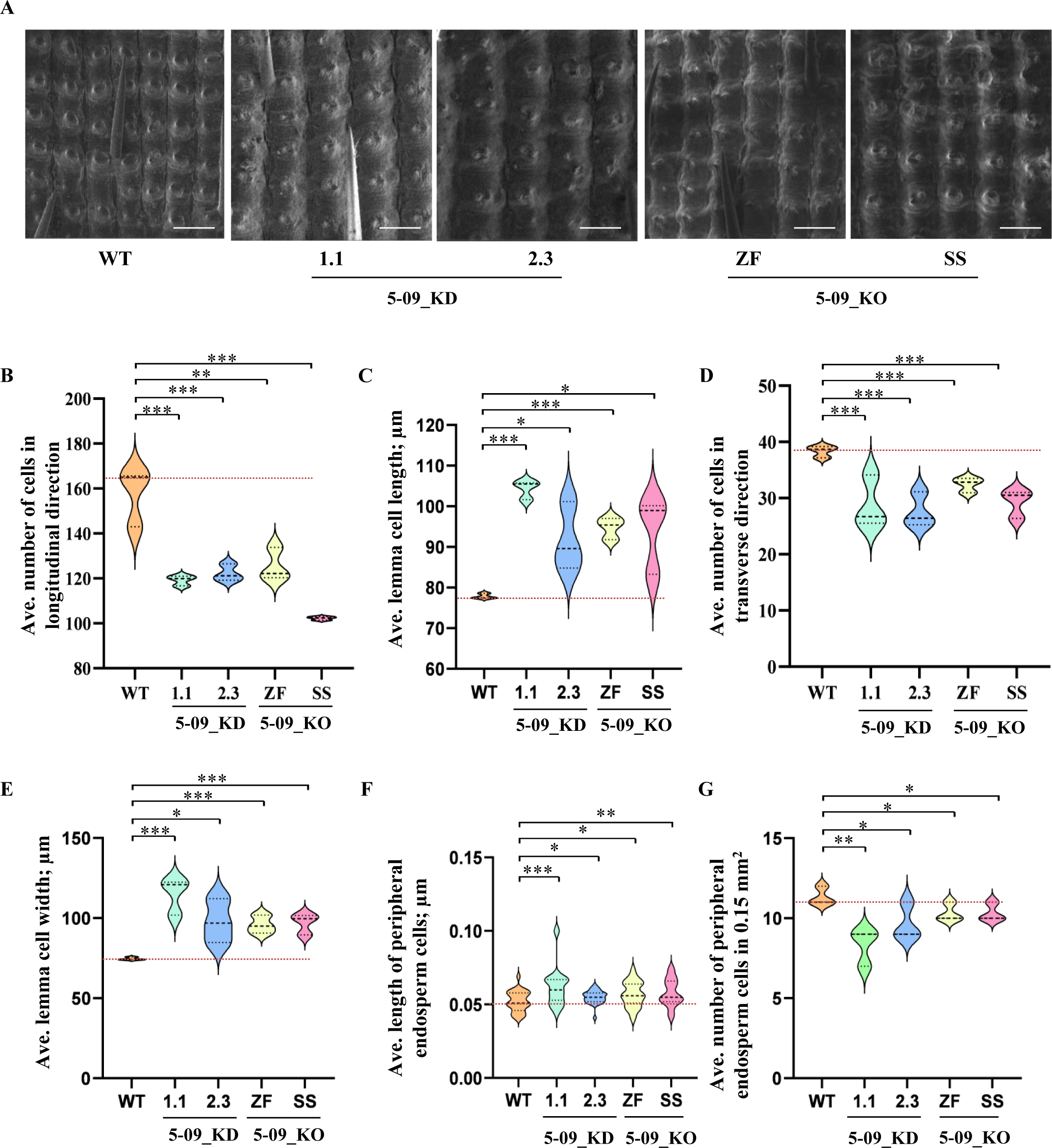
Control of grain cell size and number by *ZOS5-09.* (A) Scanning electron micrographs of the outer surface of lemma cells in transgenic spikelet hulls. Scale bars = 100 μm. Graphs show (B) average number of cells in longitudinal direction, (C) average lemma cell length, (D) average number of cells in transverse direction, and (E) average lemma cell width (n=3*15 for B-E). Graphs shows (F) average length of peripheral endosperm cells over their widest direction (n=3*9) and (G) average number of peripheral endosperm cells in a 0.15 mm^2^ area in 5-09_KD and 5-09_KO grains as compared with WT, n=3. Asterisks denote significant difference as determined by Student’s t-test (*,**,*** = p value ≤ 0.05, ≤ 0.01, ≤ 0.005 respectively). Error bars = ± SE.

**Fig. 4.**
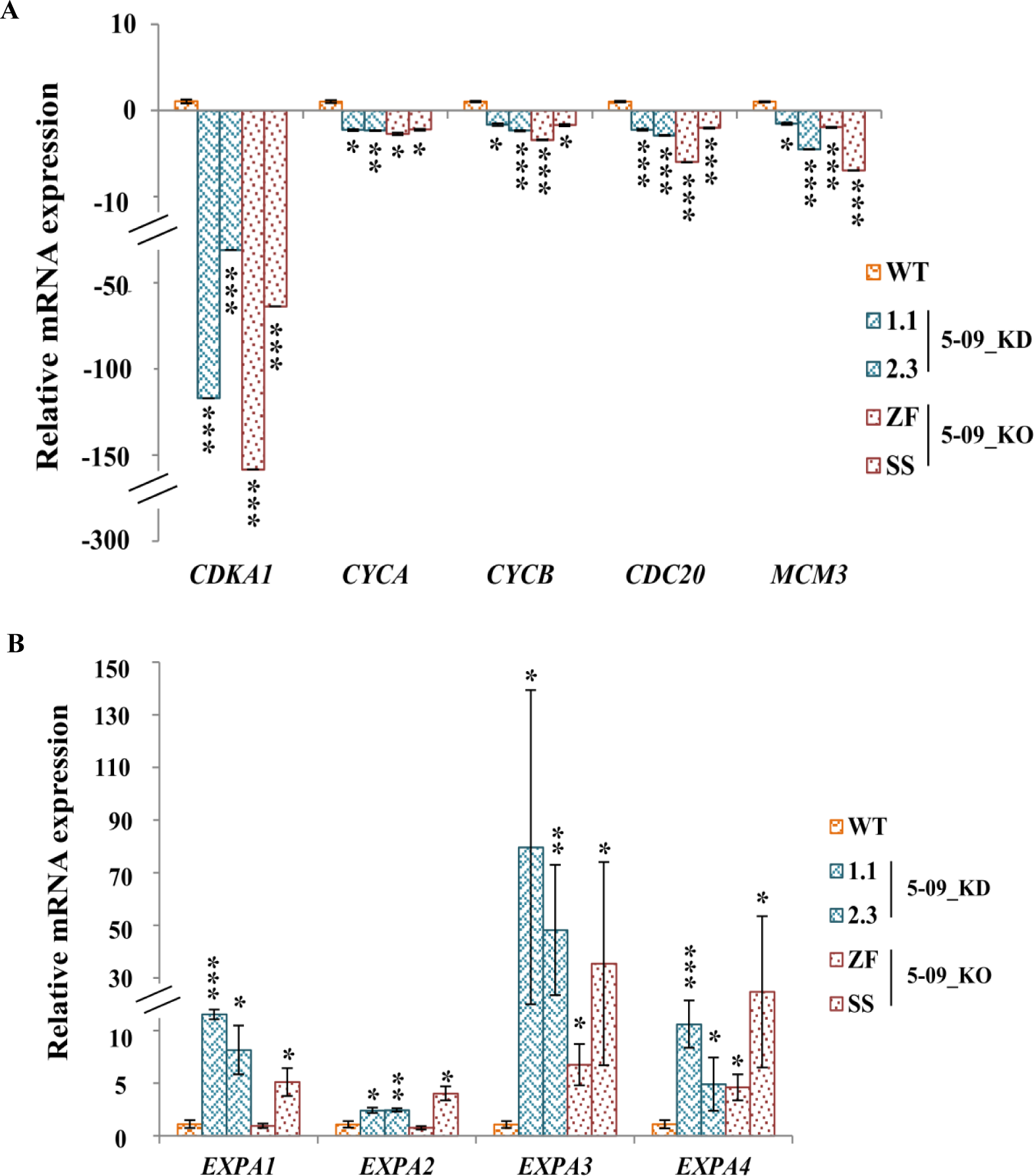
Regulation of cell division and expansion by *ZOS5-09*. Bar graphs show the relative expression of **(A)** cell cycle genes (*CDKA1, CYCA, CYCB, CDC20* and *MCM3*) and **(B)** expansin genes (*EXPA1*, *EXPA2*, *EXPA3* and *EXPA4)* in 5-09_KD and 5-09_KO grains with respect to WT grains. Asterisks denote significant differences as determined by Student’s t-test (*,**,*** = p value ≤ 0.05, ≤ 0.01, ≤ 0.005 respectively). Error bars = ± SE.

#### *ZOS5-09* affects starch and protein biosynthesis

Dehusked and transversely sectioned 5-09_KD and 5-09_KO grains had chalky patches (Fig. 5A, B). These rice grains showed a decrease in total starch (26-30%), amylose (35-49%) and amylopectin content (22-30%) (Fig. 5C) with respect to WT. The amylose: amylopectin ratio was also lower than WT in these grains indicating decreased amylose content. The chalky area showed a loss of starch granule structure and had irregularly shaped, loosely packed starch granules. However, in the translucent area, the compound granules with polyhedral sub-granules were tightly packed similar to the wild type grain (Fig. 5D). Further, qRT-PCR analysis showed a significant down regulation in the expression of all tested starch related genes in 5-09_KD and 5-09_KO with respect to wild type grains (Supplementary Fig. S15) which correlated with decreased total starch content.

**Fig. 5.**
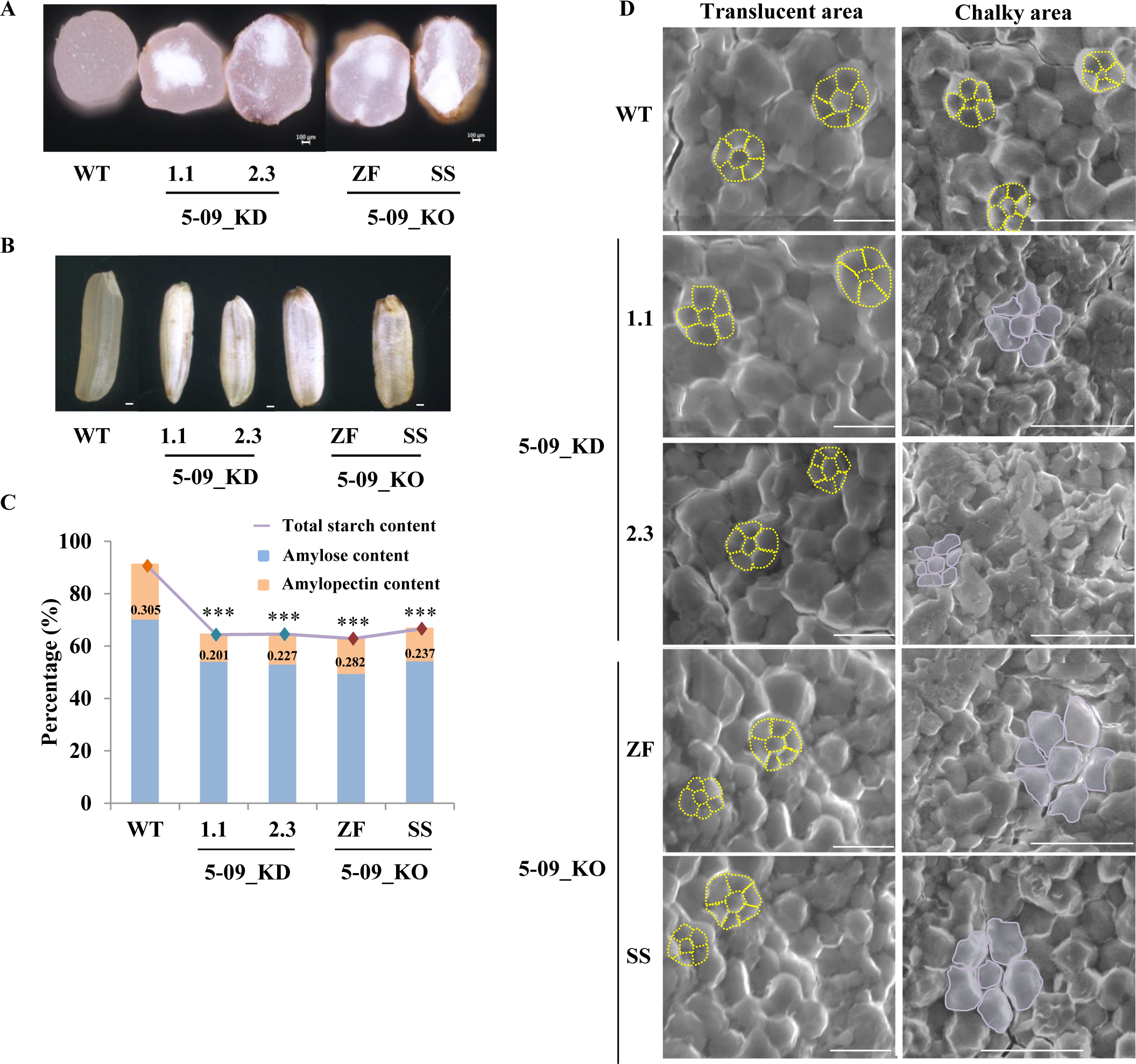
*ZOS5-09* influences starch accumulation. (A) Images show extent of grain chalkiness in transverse hand cut sections of mature grains and (B) in complete grain, of 5-09_KD and 5-09_KO plants. Scale bars = 100 μm for cut and 10 μm for complete grains. (C) Graph represents percent total starch, amylose and amylopectin content in 5-09_KD/5-09_KO and wild type matured grains. Numbers in the bars represent amylose: amylopectin ratio. Asterisks denote significant difference for total starch content as determined by Student’s t-test (*** = p value ≤ 0.005). Error bars = ± SE, n=3. (D) Scanning electron microscopy images of endosperm from mature KD/KO and WT grains, represent starch granules structure from both translucent and chalky regions. Scale bar represents 100 µm. Compound granules with polyhedral sub-granules have been denoted by yellow dashed lines and irregularly shaped, loosely packed starch granules in purple.

There was an additional decrease in the total protein content (37-41%) in 5-09_KD and 5-09_KO grains (Fig. 6A). Further, SDS-PAGE analysis of total protein from 5-09_KD and 5-09_KO grains with respect to WT grains, showed a decrease in the intensity of protein bands for glutelin precursor (∼55 kDa), glutelin acidic subunits (∼35 kDa), glutelin basic subunits (∼25kDa) and 13 kD prolamins (Fig. 6B). qRT-PCR analysis showed a decreased expression of few seed storage protein (SSP) encoding genes, *ALB3, GLB4, PRO16*, and *GLU19* (Fig. 6C). On the other hand, the expression of *ALB13*, *PRO18, GLU6* was increased in 5-09_KD and 5-09_KO grains (Fig. 6D). Hence, these data show that *ZOS5-09* knock-down/knock-out decreases total starch and protein content of rice grains, affects starch granule structure and alters the expression of certain SSP encoding genes.

**Fig. 6.**
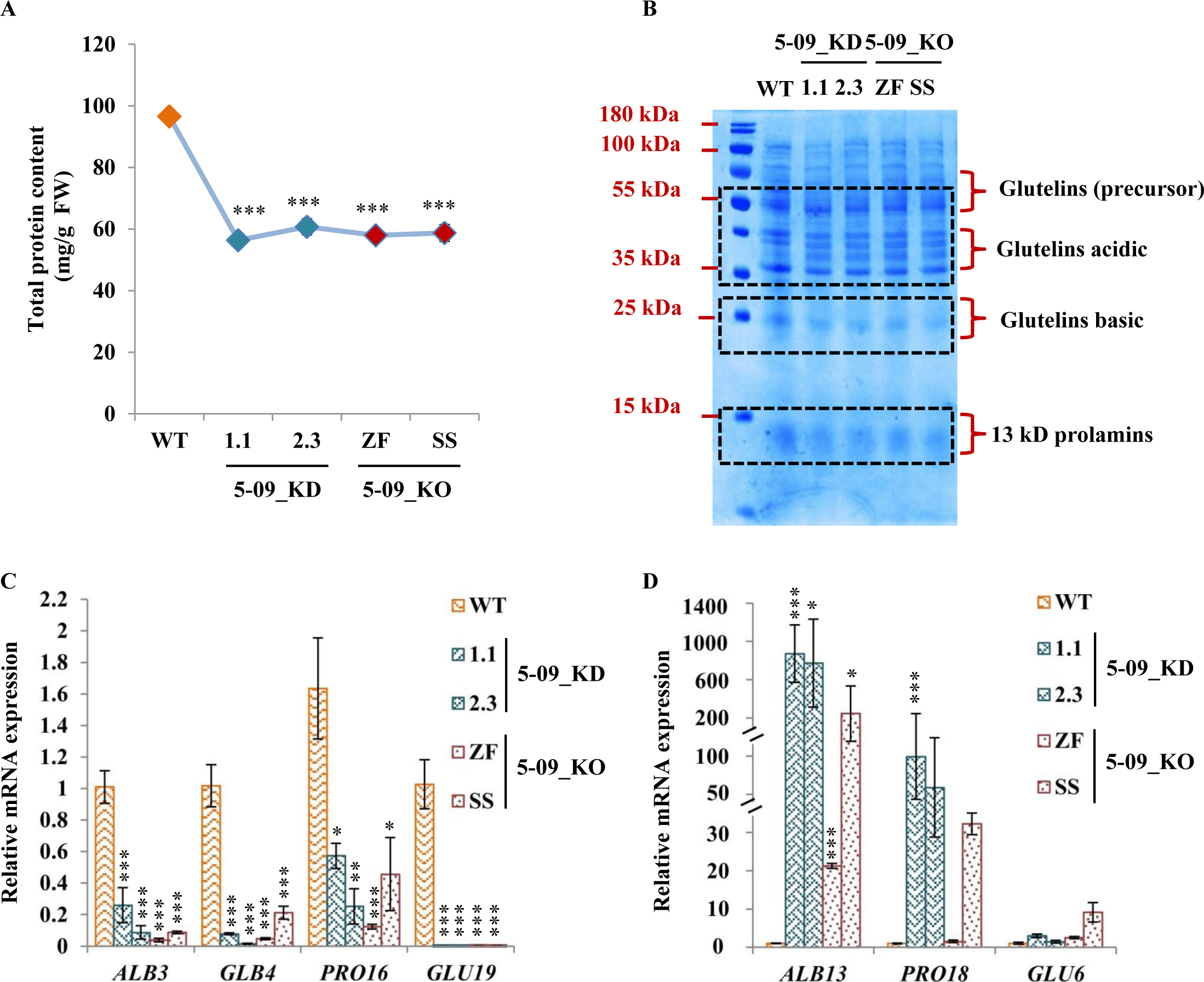
*ZOS5-09* influences total protein accumulation. (A) Total protein content in 5-09_KD, 5-09_KO and wild type S5 grains, n=3. (B) SDS-PAGE analysis of total seed protein extracted from S5 stage rice seeds of WT, 5-09_KD and 5-09_KO plants. Bands for glutelin precursor (∼50 kDa), glutelin acidic subunits (∼30 kDa), glutelin basic subunit (∼22 kDa) and 13 kDa prolamins are indicated on the right side of the SDS-PAGE gel. Black dotted boxes highlight bands with intensity variation between WT and 5-09_KD and 5-09_KO grains. Relative expression of SSP encoding genes (C) *ALB3, GLB4, PRO16, GLU19* and (D) *ALB13*, *PRO18*, *GLU6* in 5-09_KD and 5-09_KO grains with respect to WT. Asterisks denote significant difference as determined by Student’s t-test (*,**,*** = p value ≤ 0.05, 0.01, 0.005 respectively). Error bars = ± SE.

#### ZOS5-09 binds to the promoter of *GLUTELIN6* and interacts with histone deacetylase, HDA704

Since *ZOS5-09* showed the highest expression in S5 stage of seed development and altered the expression of many SSP encoding genes, it was speculated that it might be involved in SSP biosynthesis. To test the hypothesis, yeast one-hybrid (Y1H) assay was performed with promoters of *GLUTELIN6* (*pGLU6*) and *ALBUMIN13* (*pALB13*) (Supplementary Fig. S16A, B), which were upregulated in 5-09_KD and 5-09_KO grains (Fig. 6D). Growth of co-transformed yeast cells with ZOS5-09_AD and *pGLU6_pABAi* and no growth with *pALB13_pABAi* on synthetic media lacking leucine and uracil, supplemented with 200 ng/ml aureobasidin, showed the binding of ZOS5-09 to the 2 kb upstream promoter of *GLU6* but not to that of *ALB13* promoter. Deletion analysis of the promoter indicated that the binding occurred between −800 bp to −400 bp upstream to the start site (*pGLU6_4*). PLANTPAN3.0 determined the zinc finger binding site (ZF_BS) in *pGLU6_4* at −764 bp. Further, in electrophoretic mobility shift assay (EMSA) (Fig. 7A), biotin labelled 2X ZF_BS probe incubated with GST_ZOS5-09 showed a clear band shift. The binding was reduced when 50 X and 100 X unlabeled probes were used as the competitors. ZOS5-09 did not bind to mutated labeled ZF_BS. This confirmed that ZOS5-09 bound to the predicted ZF_BS of *pGLU6.* In addition, in the *in planta* reporter effector assay (Fig. 7B, C), co-transformation of effector ZOS5-09 with *pGLU6* as a reporter, repressed the expression of *GUS* gene downstream to *pGLU6*. This is in alignment with the upregulation of *GLU6* in KD and KO grains (Fig. 6D). Hence, ZOS5-09 can directly bind to the promoter of *GLU6* at the ZF_BS and repress its expression.

**Fig. 7.**
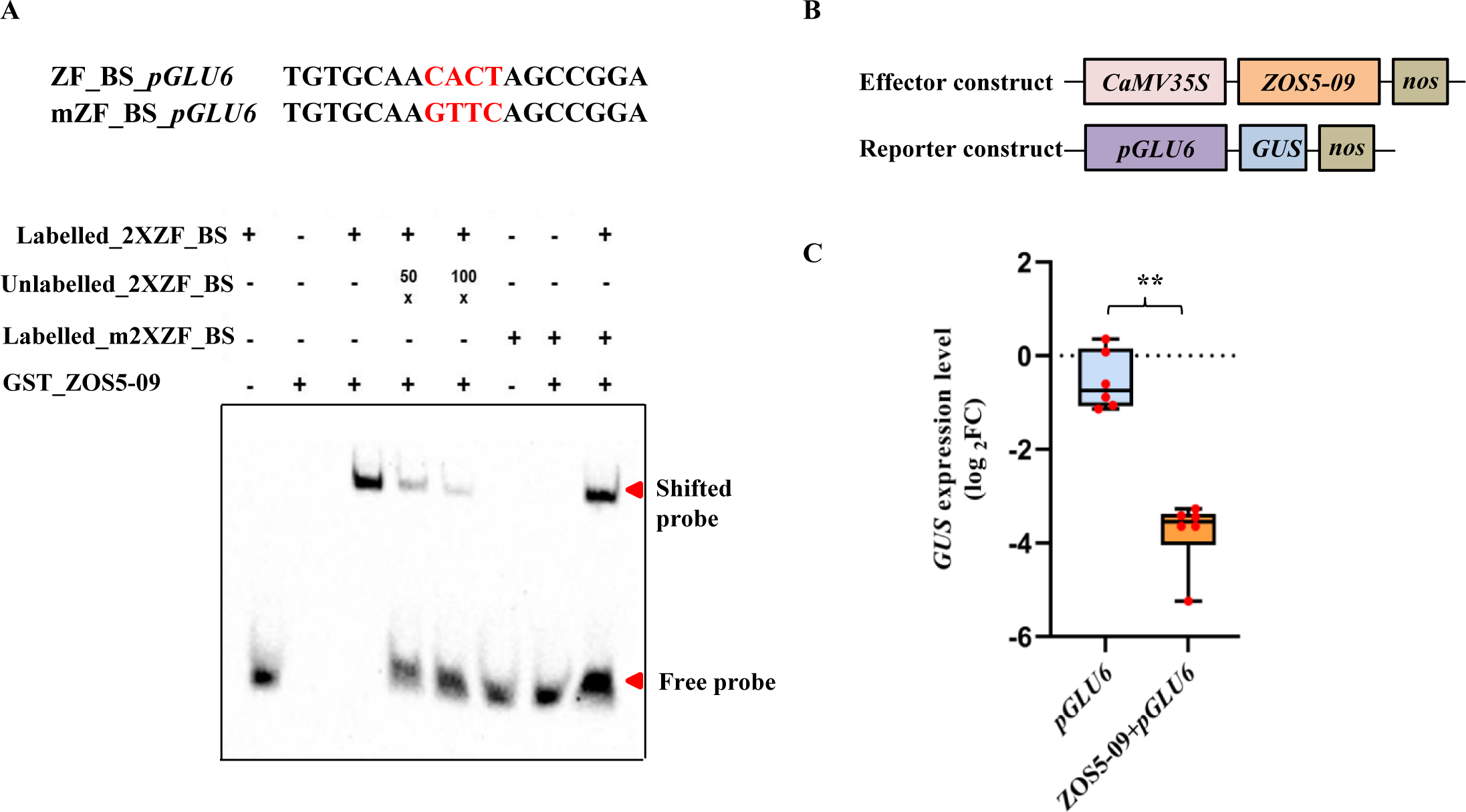
Protein-DNA interaction of ZOS5-09. (A) EMSA shows the binding of ZOS5-09 to the ZF binding site (TGTGCAACACTAGCCGGA) on *pGLU6*. GST_ZOS5-09 protein was used. Unlabelled 2XZF_BS was used as a competitor in increasing concentrations of 50X and 100X. mZF_BS indicate mutated site. (B) Schematic representation of reporter and effector constructs for reporter effector assay in *Nicotiana benthamiana.* Effector construct contains ZOS5-09 driven by *CaMV35S* promoter. Reporter construct contains *pGLU6* driving the GUS expression. (C) Reporter effector assay shows the binding of ZOS5-09 to the promoter of *GLU6.* Data are means ± SE from six biological replicates. Statistical significance for ZOS5-09+*pGLU6* was calculated with respect to *pGLU6.* Asterisks denote significant difference as determined by Student’s t test (** = p value ≤ 0.01). Error bars = ± SE.

Several reports have revealed that TFs possessing an EAR motif recruit components of the repression machinery like co-repressors and histone deacetylases (HDACs) for their function. In case of ZOS5-09, yeast two-hybrid assay and pull down assay with TOPLESS (TPL) co-repressor did not show a very clear interaction of the two, possibly indicating the role of other factors in forming the complete transcriptional complex (Supplementary Fig. S17A, B). Further, yeast two-hybrid assay of ZOS5-09 with seven HDACs (HDA704, HDA706, HDA709, HDA711, SRT701, HDT701 and HDT702) representing all three HDAC classes (Hu *et al*., 2009) (Supplementary Fig. S18, S19), resulted in growth of yeast cells co-transformed with HDA704, HDA706 and HDT701 on SD/-AHLT and gave a blue color on SD/-AHLT+X-α-gal.

This suggested that ZOS5-09 interacted with these three HDACs. Yeast two-hybrid assay of ZOS5-09_ΔDLN with HDA704 (Supplementary Fig. S3A, S19) indicated that the full length protein is responsible for interaction with HDA704. Further, in an *in vitro* pull down assay, the presence of 111.03 kDa band of MBP_HDA704 when GST_ZOS5-09 (45.14 kDa) protein was bound to beads and vice-versa, confirmed the interaction (Fig. 8A). An *in planta* co-immunoprecipitation (co-IP) assay confirmed the interaction of ZOS5-09 with HDA704 as immunoprecipitated YFP_ZOS5-09 (47.14 kDa) was detected on the immunoblot (Fig. 8B). This interaction resulted in global deacetylation as shown by the immunoblot analysis (Fig. 8C), wherein H4 histone acetylation levels in 5-09_OE calli were highly reduced in comparison with calli harboring empty vector. This meant that increased ZOS5-09 function exhibits repression through deacetylation.

**Fig. 8.**
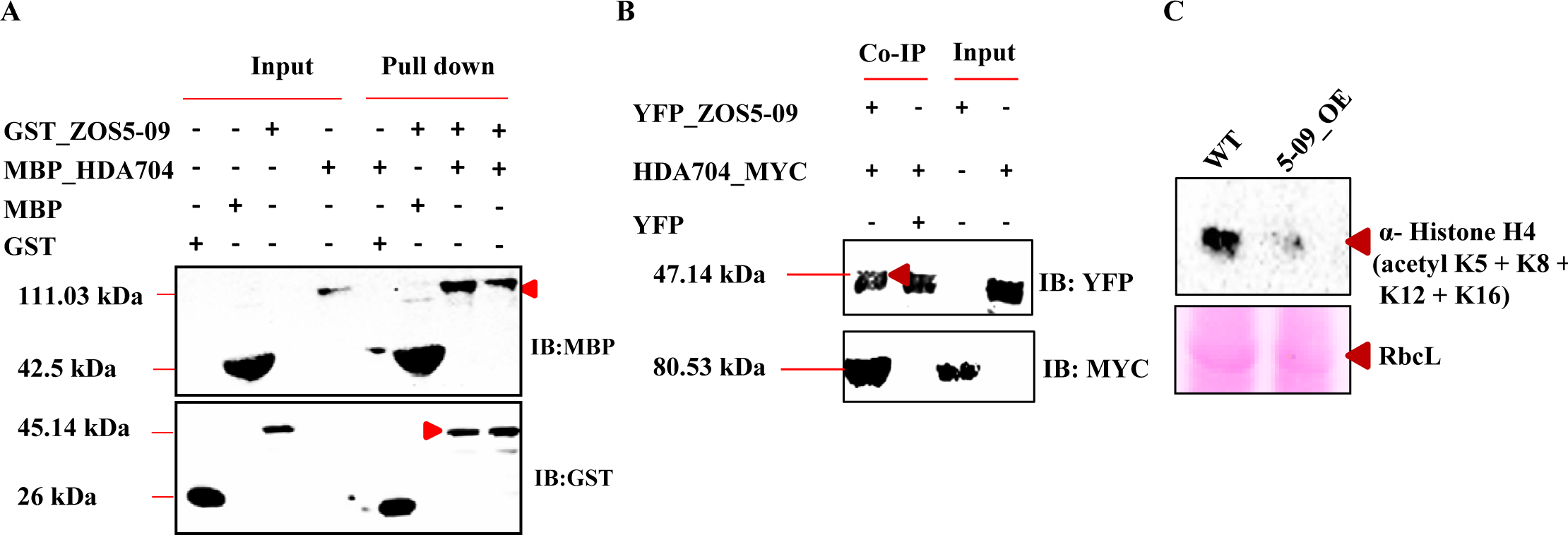
Protein-protein interactions of ZOS5-09. (A) Pull down assay shows interaction of GST_ZOS5-09 (45.14 kDa) with MBP_HDA704 (111.03 kDa). Pull down lane with the expected protein size has been indicated by a red arrow. (B) In planta co-immunoprecipitation (Co-IP) assay to confirm the interaction of YFP_ZOS5-09 (47.14 kDa) with HDA704_MYC (80.53 kDa). The red arrow indicates immunoprecipitated YFP_ZOS5-09. Input, pull down, co-IP fractions and the antibody used have been marked. IB denotes immunoblot. (C) Western blot shows the relative levels of Histone H4 (acetyl K5 + K8 + K12 + K16) in wild-type and 5-09_OE rice calli. Ponceau-S staining of Ribulose-1,5-bisphosphate carboxylase/ oxygenase large subunit (RbcL) was used as loading control.

### Discussion

C_2_H_2_ ZF transcriptional regulators are known to control developmental and stress responses in plants (Lyu and Cao, 2018, Han *et al*., 2020). In the present study, we have characterized a novel C_2_H_2_ ZF TF, ZOS5-09 that functions as a transcriptional repressor to regulate grain size and grain quality in rice. To our knowledge, this is the first such report of a C_2_H_2_ ZF TF that has a combined role in control of grain size, starch and SSP biosynthesis and filled grain number. This implies that ZOS5-09 controls all essential grain quality and quantity traits in rice. Amongst other traits, it influences plant height, leaf size and panicle number, which are important agronomic traits.

#### Control of grain size and vegetative traits by ZOS5-09

*ZOS5-09* is a seed-preferential TF, with extremely high expression in the developing rice endosperm (Fig. 1 A, Supplementary Fig. S1A, B). Based on the phenotypes of 5-09_KD and 5-09_KO plants (Fig. 2, Supplementary Fig. S11), ZOS5-09 appears to be a positive regulator of grain length and weight, filled grain number, plant height, second leaf width and panicle number. On the other hand, ZOS5-09 is a negative regulator of grain width and second leaf length (Fig. 2 D, Supplementary Fig. S11E). This shows the wider role of ZOS5-09 TF in regulating grain size and maintaining plant architecture. There are genes such as *SG2* (Yuan *et al*., 2017), *WTG1* (Huang *et al*., 2017), *RGG1* (Tao *et al*., 2020), *WG1* (Huang *et al*., 2020b) and *ZFP207* (Duan *et al*., 2021) which regulate both vegetative and grain characteristics, as observed in our 5-09_KD and 5-09_KO plant phenotypes. Grain yield is positively correlated with filled grain number per plant as well as grain size (Zhao *et al*., 2020). 5-09_KD and 5-09_KO grains exhibited reduced grain weight and filled grain number per plant (Fig. 2 E-F), which resulted in a reduced grain yield in both 5-09_KD and 5-09_KO plants. The number of unfilled grains per plant was increased in 5-09_KD and remained unaffected in 5-09_KO plants. This may be because the segmental duplicate of *ZOS5-09* was also downregulated only in 509_KD plants (Supplementary Fig. S12).

The decrease in grain length and increase in grain width (Fig 2C-D) in 509_KD and 509_KO plants is due to wider lemma cells resulting from more cell expansion in the transverse direction than the longitudinal direction. The 509_KD and 509_KO endosperm cells are also larger than WT (Fig. 3, Supplementary Fig. S13). Grain size can be regulated either through cell proliferation and/or cell expansion. Increased cell proliferation can be observed through an increase in expression of cell cycle related genes (Tsago *et al*., 2020). An increase in the expression of *EXPANSIN* genes indicates increased cell expansion (Xiong *et al*., 2023). Hence, the change in cell size and proliferation can be explained by a decrease in the expression of cell cycle related genes and increase in expression of expansins.

#### Role of ZOS5-09 in starch production

Starch accumulation is an important determinant of grain quality in cereal crops. 5-09_KD and 5-09_KO grains were chalky, had irregularly shaped, loosely packed starch granules, and had decreased total starch, amylose and amylopectin contents (Fig. 5 A-D). The amylose: amylopectin ratio was also reduced. These grains showed decreased expression of all eight tested starch related genes (*GBSS1*, *AGPS1*, *AGPL1*, *SSIII*, *SSI*, *FLO7*, *BeIIb*, *MST4*) (Supplementary Fig. S15). Of these, *GBSSI* is responsible for amylose synthesis (Pérez *et al*., 2019) and *AGPL1* and *AGPS1* are involved in starch biosynthesis (Meng *et al*., 2020). *SSI*, *SSIII* and *BeIIb* are associated with accumulation of amylopectins (Huang *et al*., 2016, Okpala *et al*., 2022). *FLO7* is essential for proper amyloplast development (Zhang *et al*., 2015). *MST4* encodes for a monosaccharide transporter (Wang *et al*., 2007) and its low expression indicates reduced accumulation of sugars in sink tissues in 5-09_KD and 5-09_KO grains. Our results suggest KD and KO of ZOS5-09 affected total starch accumulation and proper amyloplast development, possibly due to decreased expression of many starch related genes.

#### The direct effect of ZOS5-09 on SSP biosynthesis

Accumulation of SSPs is another determining factor for grain quality in cereal crops. We report that 5-09_KD and 5-09_KO grains showed reduced total protein content (Fig. 6A-B) and decreased expression of four SSP encoding genes (*ALB3*, *GLB4*, *PRO16*, and *GLU19*) from rice (Fig. 6C). These grains also showed increased accumulation of transcripts of three *SSPs* (*ALB13, PRO18, GLU6*) (Fig. 6D). This is possible because often there is a balance between different SSPs (Kawakatsu *et al*., 2010). Similar to our data, where both starch and protein content are affected, *opaque3* mutants have decreased protein content and reduced starch accumulation in rice endosperm (Cao *et al*., 2022). To further support the direct regulation of SSPs by ZOS5-09, we have shown that ZOS5-09 binds to the ZF_BS in the promoter of *GLU6* (Fig. 7A) and represses it *in planta* (Fig. 7B-C). In accordance, *GLU6* is upregulated in 5-09_KD and 5-09_KO grains, which have decreased expression of *ZOS5-09* (Supplementary Fig. S11C). Only a few TFs repressing the expression of SSP encoding genes are known, such as TaNAC100 on *GLU-1* (Li *et al*., 2021), TaNAC019 on *glutenin* (*HMW-GS*) (Gao *et al*., 2021) and OsNAC20 and OsNAC26 on *GluA1*, *GluB4*, *α-globulin*, and *16 kD Prolamin* (Wang *et al*., 2020b), and ZOS5-09 adds to our knowledge of the same.

#### ZOS5-09 as a transcriptional repressor

ZOS5-09 has a DLN motif (Agarwal *et al*., 2007) which is able to successfully repress reporter gene activity in yeast (Singh *et al*., 2019). The *in planta* experiments detailed in this article (Supplementary Fig. S3C, Fig. 7B-C), further strengthen our previous data. Repressors are known to interact with histone deacetylases and co-repressors (Zhuang *et al*., 2020). ZOS5-09 interacts with a histone deacetylase, HDA704 (Fig. 8A-B) and causes global deacetylation (Fig. 8C). HDA704 is a H4 type deacetylase of RPD3/HDA1 family. It functions by decreasing the H4 acetylation levels in the downstream target genes (Zhao *et al*., 2021, Yoon *et al*., 2022). Downregulation of HDA704 results in reduced plant height and altered leaf morphology (Hu *et al*., 2009). These phenotypic results are similar to the 5-09_KD and 5-09_KO phenotypes and indicate that a decreased activity of ZOS5-09, which acts in conjunction with HDA704, negatively affects plant height and leaf width.

#### The essentiality of NoRS and an equilibrated expression of *ZOS5-09*

The ectopic or seed-preferential overexpression of *ZOS5-09* (Supplementary Fig. S4, S5B, D) was lethal. The seed-preferential overexpression of ZOS5-09 with the NoRS deleted (Supplementary Fig. S5 C, E-P), did not result in any grain filling. In view of these data and the grain phenotypes of 5-09_KD and 5-09_KO plants, we speculate that a strictly regulated expression of this gene is essential, the mechanism of which needs further exploration. This implies that ZOS5-09 needs to be present in controlled amounts in the plant for it to function in a beneficial manner. We have previously observed that ectopic overexpression of seed-preferential *ONAC025*/*SS1* is also detrimental (Mathew *et al*., 2020) because of an expansion of the reproductive transcriptome into the vegetative zones, which may be the case here as well. ZOS5-09 has a C-terminal NoRS (Fig. 1B). Our conclusion is that the NoRS is essential for the function of ZOS5-09 as its deletion by CRISPR/Cas9 (SS data in Fig. 2-6) imparts the same phenotype as when the complete protein is altered (ZF data in Fig. 2-6) or the gene is knocked down (KD data in Fig. 2-6). Though this may need further examination, our study is the first such report about the functional role of NoRS, as per literature survey.

#### Conclusion

Since *ZOS5-09* functions as an important regulator of grain size, starch and SSP synthesis, this study helps in uncovering the co-regulated mechanism of size along with starch and SSP accumulation in rice seeds. We speculate that reduced plant height and grain size in 5-09_KD and 5-09_KO plants might lead to reduced demand for starch and protein biosynthesis, which eventually affects grain quality (Fu and Xue, 2010). Our study concludes that a stable equilibrated expression of *ZOS5-09* is required for proper plant growth and development. Based on our findings, we propose a working model of ZOS5-09 in regulating rice development (Fig. 9). ZOS5-09 interacts with HDA704 and acts as a repressor by global deacetylation. Ectopic overexpression and downregulation/ knock-out, both result in strong phenotypes and affect plant growth. Knock-down or knock-out of *ZOS5-09* reduces plant height, second leaf width and panicle number. In terms of grain phenotype, these plants have decreased grain length, grain weight, filled grain number and increased grain width. The data that ZOS5-09 binds to ZF_BS (Fig. 7A) hints at the fact that other promoters with ZF_BS might be controlled by this TF and identification of other such genes will explain the KD and KO phenotypes. Since *ZOS5-09* promoter directs endosperm-specific expression and ZOS5-09 binds directly to *GLU6* promoter, it has an irreplaceable role in rice grain filling. These findings provide novel insights about the function of *ZOS5-09* and will aid in understanding the regulation of grain size and grain quality.

**Fig. 9.**
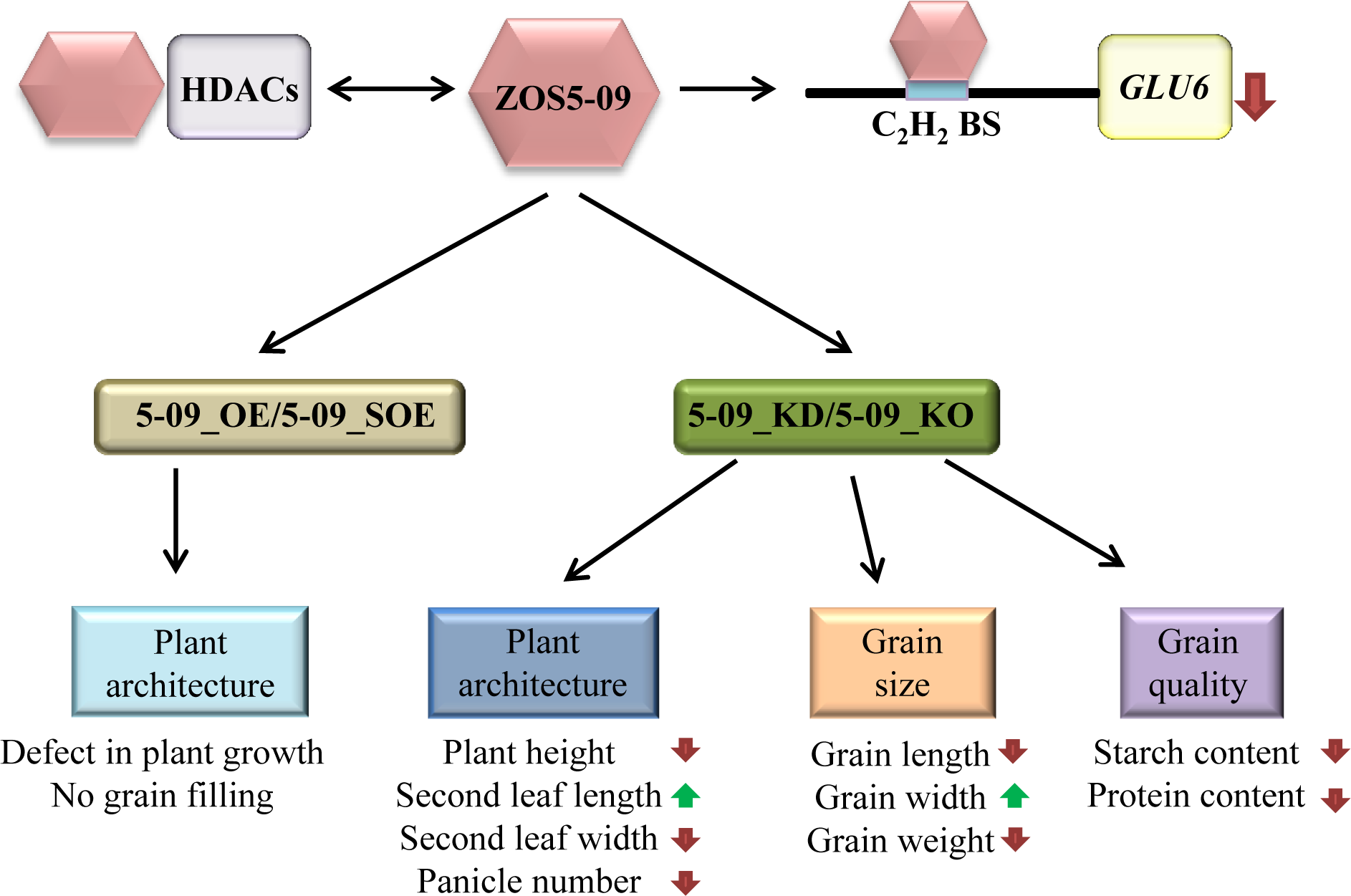
Working model of *ZOS5-09*. The proposed model depicting the role of *ZOS5-09* in regulating plant architecture, grain size and grain quality in rice. ZOS5-09 (indicated by peach pink hexagon) interacts with histone deacetylases (indicated in light purple) and binds to C_2_H_2_ binding site (indicated by blue colour) present in *GLU6* promoter and represses its expression through deacetylation. 5-09_OE plants show a defect in plant growth and development. Suppressed expression or knock-out of *ZOS5-09* negatively affects plant architecture (plant height, leaf length, leaf width, and number of panicles per plant), grain size (grain length, width, and weight) and grain quality (starch and protein accumulation). The relationship between plant phenotype and *GLU6* expression is speculative and will require further investigation to determine other downstream genes contributing to the phenotype. Upward arrow show increased phenotype and downward arrow showed decreased expression or decreased phenotype.

### Experimental procedures Plant materials

Rice (*Oryza sativa* L. cv. IET-10364) was grown in NIPGR fields, New Delhi, India under natural conditions during rice growing season. For expression analysis of *ZOS5-09*, early panicle developmental stages, P1 (0-5 cm) and P2 (5-10 cm), and seed developmental stages (S1-S5) (Sharma *et al*., 2012), flag leaf, second leaf, leaf sheath, root, callus, node and internode, were harvested using liquid nitrogen and stored at −80°C for experimental analysis.

### RNA extraction and qPCR analysis

Total RNA from S1-S5 seed stages was isolated using modified GH protocol. Total RNA was isolated using extraction buffer (50 mM Tris-HCl pH 9.0, 20 mM EDTA pH 8.0, 150 mM NaCl, 1% N-lauryl sarcosyl (sodium salt) and 5 mM DTT) and GH buffer [8 M guanidine hydrochloride, 20 mM 2-(N-Morpholino) ethanesulfonic acid}, 0.5 M EDTA, 50 mM β-mercaptoethanol) and RNA was precipitated using 3 M sodium acetate (pH 5.2) and chilled ethanol. TRIzol Reagent^®^ Solution (Invitrogen^TM^; USA) was used to isolate RNA from other rice tissues, i.e., node, internode, flag leaf, root and panicle stages. RNase-free DNase Set (Qiagen; Germany) was used to remove genomic DNA contamination, according to manufacturer’s instructions. RNA was purified using RNeasy^®^ MinElute Cleanup Kit (Qiagen; Germany). The concentration and purity of RNA samples were checked by NANODROP 2000c Spectrophotometer (Thermo Scientific; USA). RNA samples were also checked by running on 1% denaturing agarose gel in 1X MOPS buffer (400 mM MOPS (3-(N-morpholino) propanesulfonic acid), 99.6 mM sodium acetate, 20 mM EDTA (Ethylenediamine tetraacetic acid)) with 1.1 % formaldehyde. 2 μg of total RNA was used to synthesize cDNA using the High-capacity cDNA Reverse Transcription kit (Applied Biosystems; USA). For a single reaction, 2 µl of 2X RT buffer, 0.8 µl 25X dNTP mix, 2 µl of 10X RT random primers, 1 µl of reverse transcriptase, 2 µg of RNA was added and volume was set to 20 μl with MQ water. PCR cycle for preparation of cDNA was set up as, 25 °C for 10 min, followed by 40 cycles of 37 °C for 40 mins, and a cycle of 85 °C for 5 min. cDNA was then stored at −80°C. Quantitative real time PCR was performed on 7500 Fast Real-Time System (Applied Biosystems; USA) using SYBR Green (Applied Biosystems, USA) and 7500 software v2.0.1 was used for data analysis with three biological and three technical replicates. Primers for qPCR were designed using Primer Express^TM^ software version 3.0 and amplicon length range was 60-120 nt long. The specificity of primers was checked using nucleotide BLAST program. For qPCR, 1 μl of 1:10 diluted cDNA was mixed with 5 μl SYBR Green and 0.9 μl each of 10 μM forward and reverse primer, volume of the reaction was made up to 25 µl with MQ water. The qPCR cycle was set as: 50 °C for 2 min, 95 °C for 2 min, followed by 40 cycles of 95 °C for 15 s, 60 °C for 60 s, and a cycle of 95 °C for 15 s, 60 °C for 15 s, and 95 °C for 15 s followed by melting curve analysis. To find the relative expression of target genes, qPCR was performed at S5 stage of transgenic grains with respect to wild type of same stage. The expression level of three reference genes, Actin (*ACT1*), Ubiquitin (*UBQ5*) and GAPDH (*GAPDH*) were compared. The relative expression of *UBQ5* and *GAPDH* gene was calculated with respect to *ACT1* as reference genes. *UBQ5* and *GAPDH* expression did not change significantly (Table S2) in different transgenic materials, so we used *ACT1* as the endogenous control for normalization for all qPCR experiments in this study. Fold change was calculated by comparative C_T_ (ΔΔC_T_) method (Rao *et al*., 2013). For expression analysis of *ZOS5-09* in developmental stages, fold change was calculated using a constant C_T_ (= 30) value (Mathew *et al*., 2016). This value is considered as no expression. The C_T_ value of no template control in all reactions was undetermined. All the essential information under the MIQE checklist, used in the experiment, has been mentioned above.

### Promoter analysis of *ZOS5-09* and generation of rice transgenic plants

A 1618 bp region of *ZOS5-09* promoter was used for promoter analysis. The promoter sequence was scanned for presence of *cis*-acting elements using PLACE and PLANTCARE databases. Further, it was amplified from genomic DNA of *Oryza sativa* var. IR64 and cloned into pMDC164 and pMDC110 (Curtis and Grossniklaus, 2003) to produce *pZOS5-09::GUS* and *pZOS5-09::GFP,* respectively, through the entry vector, pENTR^TM^ D-TOPO^®^ (Invitrogen^TM^, USA) using Gateway^®^ based cloning (Invitrogen^TM^, USA). The primers used in this study are listed in Table S1. Rice seeds were transformed with *Agrobacterium* strain EHA105 carrying the fusion constructs (*pZOS5-09::GUS* and *pZOS5-09::GFP)* using a scutellum transformation based protocol (Toki *et al*., 2006). For generation of transgenic plants, rice seeds were sterilized using 0.1% HgCl_2_ and 2-3 drops of teepol and inoculated on NB media (media contains N6 major (Stock I), B5 minor (Stock II), Fe-EDTA (Stock III), and organic source (Stock IV). 1 L of N6B5 media was prepared by adding 50 ml stock I, 1 ml stock II, 10 ml stock III, 1 ml stock IV, 2.878 g L-proline, 500 mg L-glutamine, 300 mg CEH (casein enzyme hydrolysate), 30 g of sucrose, and 4 g phytagel per liter of solution with a pH-5.8, supplemented with 2 mg/ml of 2,4-D (2,4-dichlorophenoxyacetic acid)) for callus induction at 32 °C with continuous light for 5 days. After the incubation period, primary culture of *Agrobacterium* harboring *pZOS5-09::GUS* or *pZOS5-09::GFP* construct was co-cultivated with rice embryogenic calli and these were placed on N6B5 medium (with 2 mg/ml of 2, 4-D, 100 μM acetosyringone, 4 g phytagel per liter of solution, pH-5.2). The plates containing co-cultivated calli were incubated for two days at 28 °C in dark conditions. After incubation, the co-cultivated calli were transferred to the selection media (N6B5 media with pH-5.8, supplemented with 300 mg/L of augmentin and 50 μg/ml of hygromycin). The selection plates were further incubated at 32 °C for 28 days in continuous light. Proliferating calli generated during selection were then transferred onto regeneration medium (50 ml stock I, 1 ml stock II, 10 ml stock III, 1 ml stock IV, 300 mg CEH, 30 g of sucrose, 4 g phytagel per liter of solution, 300 mg/ml of augmentin, 1 mg/l of BAP and 50 mg/l of hygromycin, pH-5.8) and were incubated at 28 °C with 12:12 light and dark photoperiod conditions (12 h light/12 h dark; PPFD (Photosynthetic Photon Flux Density) 350 µmol m^−2^ s^−1^) till regeneration. After the shoots attained a length of 2–3 cm, they were transferred to the rooting media (25 ml stock I, 500 μl stock II, 5 ml stock III, 500 μl stock IV, 20 g of sucrose, 2 g phytagel per liter of solution, 300 mg/ml of augmentin and 50 mg/l of hygromycin with a pH-5.8) and the tubes were incubated at 28 °C with 12:12 light and dark photoperiod conditions. Plantlets were grown hydroponically using Yoshida medium (Yoshida *et al*., 1971) and were transferred to pots in contained greenhouse conditions at 16-h-light/8-h-dark cycle (PPFD 250 µmol m^−2^ s^−1^) at 28 °C.

### Analysis of GUS and GFP activity

A histochemical assay was performed to check the expression of *GUS* reporter gene under the control of *ZOS5-09* promoter (Jefferson *et al*., 1987). Longitudinal hand-cut sections of seed at stages S1-S5 and control tissues (panicle, root, leaf) were incubated in GUS buffer (0.5 mM potassium ferricyanide, 0.5 mM potassium ferrocyanide, 0.1% (v/v) Triton-X-100, 50 mM phosphate buffer (pH 7.0), 10 mM sodium EDTA and 1 mg/ml (w/v) X-gluc (5-bromo-4-chloro-3-indolyl-β-D-glucuronide)) containing 10% methanol at 37 °C for 16-20 h. Hand cut sections of seeds at various stages of development were also visualized. GUS activity indicated by blue colour was visualized under Nikon AZ100 microscope. For GFP analysis, hand-cut sections of seed at stages S1-S5 and control tissues (panicle, root, leaf) were observed under fluorescence microscope (LP515, Leica Gmbh, Germany). The microscope was fitted with digital camera, DC500, to capture the photographs (Leica Gmbh, Germany).

### Subcellular localization

Full-length coding sequences of *ZOS5-09* (174 amino acids) and *ZOS5-09* with nucleolar retention signal removed (168 amino acids) were amplified and cloned in pSITE-3CA vector (Chakrabarty *et al*., 2007), in-frame with YFP tag under the control of the *CaMV*-35S promoter through the intermediate vector, pENTR^TM^ D-TOPO^®^ (Invitrogen^TM^, USA). pENTR constructs were proceeded for LR reaction and the insert was transferred into the destination vector, pSITE-3CA vector to produce YFP-5-09 and YFP-(ΔNoRS)-5-09 using Gateway^TM^ LR Clonase^TM^ II Enzyme mix (Invitrogen^TM^; USA). Fusion YFP constructs, i.e., YFP-5-09, YFP-(ΔNoRS)-5-09 and empty YFP were transiently expressed into onion epidermal cells through biolistic bombardment using PDS-1000 particle delivery system (BioRad, USA) (Scott *et al*., 1999) and in three-week-old leaves of *Nicotiana benthamiana* using *Agrobacterium*-mediated infiltration (Norkunas *et al*., 2018). YFP signal was visualized at 514 nm under TCS-SP5, confocal laser microscopy (Leica, Germany). DAPI (4′,6-diamidino-2-phenylindole) fluorescence and P_R82_::2xRFP (Huang *et al*., 2014), used for nucleus staining, were observed at 358 nm and at 588 nm, respectively.

### Trans-activation assay of ZOS5-09

Coding sequence of ZOS5-09 was cloned in pGBKT7 vector (Takara™ Bio USA, Inc), forming ZOS5-09_pGBKT7 construct. The construct was transformed into yeast strain AH109 using EZ-Yeast^TM^ Transformation kit (MP Biomedicals; USA) and plated on SD/-Trp. The transformed colonies were patched on triple drop-out (TDO) which contains synthetic media lacking adenine, tryptophan and histidine (SD/-Trp/-His/-Ade) and on the synthetic media lacking tryptophan SD/-Trp with 80 mg/L X-β-gal (5-bromo-4-chloro-3-indolyl-β-D-galactopyranoside). rGAL4 and empty pGBKT7 vectors were used as positive and negative control.

### Preparation of overexpression, knock-down and knock-out constructs

For generation of *ZOS5-09* (LOC_Os05g38600) overexpression plants (5-09_OE), 594 bp sequence was amplified from full length cDNA of *Oryza sativa* subsp. *indica* cv. IR64. The amplified sequence of *ZOS5-09* was cloned in pB4NU vector (Borah and Khurana, 2018) between *Bam*HI and *Sac*I sites, downstream to the *Ubiquitin* promoter. Seed-preferential overexpression constructs (5-09_SOE_1 and 5-09_SOE_2) were generated by cloning in pG6SOE vector downstream to the *GluD-1* promoter using Gateway^®^ based cloning (ThermoScientific^TM^, USA) as shown in Supplementary Fig. S5A. 5-09_SOE_1 contain 525 bp full length coding sequence and 5-09_SOE_2 contains 507 bp coding sequence without NoRS sequence as shown in Supplementary Fig. S5B-C. To generate *ZOS5-09* RNAi (5-09_KD) construct, 565 bp sequence of *ZOS5-09* along with CACC were amplified and cloned into Gateway^®^ entry vector pENTR^TM^/D-TOPO^®^ (ThermoScientific^TM^, USA). The gene was then cloned in pANDA vector (Reynolds *et al*., 2004) in an opposite orientation in between two *attR* sites, using Gateway^®^ based cloning (ThermoScientific^TM^, USA). To design *ZOS5-09* knock-out mutants (5-09_KO) by CRISPR analysis, a single gRNA targeting the first ZF domain and another gRNA targeting nucleolar retention signal (SS) were designed from coding region of *ZOS5-09* to generate ZF-5-09_KO and SS-5-09_KO constructs using CRISPR-GE tool (http://skl.scau.edu.cn/targetdesign/) (Xie *et al*., 2017). ZF_sgRNA was positioned 3 bp downstream to the start codon and SS_sgRNA was positioned at 473 bp after the start codon. This unique stretch of gRNA along with the tRNA sequence was cloned in pRGEB32 vector (Addgene, USA) using a standard protocol (Xie *et al*., 2015). gRNA-tRNA sequence was amplified using pGTR vector (Addgene, USA). The amplification of gRNA-tRNA sequence involved two standard PCR reactions and one over lapping PCR reaction. First reaction involved the amplification of gRNA-tRNA sequence (127 bp) using L5AD5 forward primer and gRNA reverse primer. The second reaction involved the amplification of gRNA-tRNA sequence using gRNA forward primer and L3AD5 reverse primer (127 bp). Following this, an overlapping PCR reaction was carried out which involved two reactions, the former reaction involved annealing of 20 ng of elute each from first and second reaction. The annealing reaction was stopped after 10 cycles and second reaction mixture (containing same PCR components with 5 μM L5AD5 forward and L3AD5 reverse primer) was added and reaction continued up to 20 cycles. The reaction cycle yielded a 244 bp fragment which was then digested with *Fok*I enzyme. *Fok*I digested gRNA-tRNA sequence was then cloned in the pRGEB32 vector using Golden Gate kit (ThermoScientific^TM^, USA). All the primer sequences are listed in Table S1. The positive clones for 5-09_OE, 5-09_KD and 5-09_KO construct were confirmed by sequencing. The positive constructs were introduced into EHA 105 *Agrobacterium* strain and transformed into *Oryza sativa* subsp. *indica* cv. IET-10364 using a scutellum-transformation based protocol (Toki *et al*., 2006). 50 μg mL^−1^ hygromycin was used for selection of transgenic calli.

### Screening of rice transgenic plants and identification of mutation in *ZOS5-09* knock-out lines

5-09_OE calli were screened for *GUS* expression through histochemical *GUS* staining (Jefferson *et al*., 1987). In addition, 5-09_OE, 5-09_KD and 5-09_KO plants were screened for the presence of *HptII* gene (850 bp) using their genomic DNA as a template for amplification. Mutations in 5-09_KO lines in T_0_ generation were identified by amplification of the target region using primers flanking the target region of *ZOS5-09*. The amplified target sequence was cloned in pJET vector using CloneJET PCR Cloning Kit (ThermoScientific^TM^, USA). Eight positive clones from each knock-out line were used for Sanger sequencing. The raw sequences were aligned to *ZOS5-09* CDS using Clustal Omega tools (https://www.ebi.ac.uk/Tools/msa/clustalo/). In the subsequent generation, plants were screened for the presence of *HptII* gene (850 bp), *Cas9* (549 bp) and target (321 bp using U3 promoter specific primers). All the primers are listed in Table S1. Off-target genes were selected using CRISPR-GE tool (http://skl.scau.edu.cn/targetdesign/) (Xie *et al*., 2017) on the basis of high off-target score. Target sequence of off-target genes was amplified with gene-specific flanking primers using genomic DNA as a template from all plants. The amplified product was cloned in pJET vector using CloneJET PCR Cloning Kit (ThermoScientific^TM^, USA). Eight positive clones from each plant were given for sequencing. All the primers are listed in Table S1.

### Phenotypic analysis of transgenic plants

ZOS5-09 seed preferential overexpression, knock-down, knock-out as well as wild type (WT) plants were used to study vegetative and grain traits. Various vegetative characters like plant height, number of tillers, stem width, flag leaf angle, flag leaf length and width, second leaf length and width, number of panicles and panicle length were measured using flexible inch-tape when plants were completely matured. Three biological replicates from each line, were used for analysis. Grains from each plants were then harvested and dried at 42°C overnight. Grain length and width were measured using WinSEEDLE^TM^ software (Regent instrument Inc.; Canada). Grain weight was measured using analytical balance. Three biological replicates from each line, with 50 grains per plant, were used to measure change in grain length, grain width and grain weight. Images of plants and seeds were photographed using a Nikon-5200 DSLR camera.

### Scanning electron microscopy and grain chalkiness

For the morphological characterization of rice seed husk, central lemma cells of intact mature seeds were scanned under scanning electron microscopy (SEM, Zeiss, Germany). Images were captured at 300X magnification. To observe change in endosperm cells, dehusked mature grains of 5-09_KD and 5-09_KO with WT grains were cut transversely with scalpel, thin sections were stained with 0.25% Toluidine blue-O/TBO and images were observed under light microscope (Eclipse 80i, Nikon; Japan) at 4X magnification. The measurements of outer epidermal cells of lemma and endosperm cells were done using ImageJ software (Schneider *et al*., 2012). To study morphology of starch granules, 5-09_KD and 5-09_KO mature grains, along with WT, were dehusked manually and seeds were cut in a transverse manner using a sharp scalpel. The central portion of the endosperm was scanned and images were captured at 2500X. To study grain chalkiness, the mature grains of WT and 5-09_KD and 5-09_KO were manually given a transverse cut and observed under stereozoom microscope (AZ100, Nikon; Japan) at 2.5X magnification.

### Estimation of starch content in rice grains

100 mg dehusked mature grains of 5-09_KD, 5-09_KO and WT were used to determine total starch content, using total starch assay kit (AA/AMG, K-TSTA; Megazyme) according to the manufacturer’s instructions and the absorbance was measured at 510 nm using spectrophotometer (BioRad, USA). 100 mg dehusked mature rice seeds were used to measure total amylose content by iodine binding affinity (Juliano, 1971). Seeds were ground into a fine powder in a mortar-pestle with the help of liquid nitrogen. The fine powder was transferred into a 100 ml volumetric flask, and 1 ml of 95% ethanol and 9 ml of 1 M NaOH were added. The solution was boiled for 30 mins at 100 °C to gelatinize starch. Then, the flask was kept at room temperature for 10 mins and adequate distilled water was added to make up the volume to 100 ml. 5 ml of this starch solution was added to another 100 ml volumetric flask and 1 ml of 1 M acetic acid and 2 ml of Lugol’s solution (0.2 gm of iodine and 2 gm of potassium iodide in 100 ml MQ) were added into it and mixed well. The solution was left to rest for 30 mins. Following this, a calorimetric assay was performed by taking the absorbance at 620 nm in the spectrophotometer (BioRad, USA). The amylose content was calculated based on the standard graph, amylose content (%)=0.682136+47.5978×Abs620 (Avaro *et al*., 2009). The amylopectin content was calculated by subtracting percent amylose content from percent total starch content.

### Total protein isolation from rice seeds

Using liquid nitrogen, 100 mg mature rice seeds from S5 stage were ground to a fine powder in a mortar and pestle. Total seed protein was extracted using 1 ml of protein extraction buffer (20 mM Tris, pH 7.5, 150 mM NaCl, 1 mM EDTA pH 8, 1 mM DTT, 1 mM PMSF) (Lang *et al*., 2013). Following this, the mixture was centrifuged at 13,000 rpm for 15 minutes. The supernatant contained total protein and was taken to a fresh tube. Total protein concentration of each sample was identified using Bradford assay (Bradford, 1976). Absorbance of the protein sample was measured at 595 nm and 40 µg of the total protein from each sample, were then separated by 10% SDS-PAGE (Kresge *et al*., 2006).

### Yeast one-hybrid assays

For prey construct, the full-length coding sequence of *ZOS5-09* was PCR amplified and cloned in pGADT7 vector (Takara™ Bio USA, Inc) to produce ZOS5-09_pGADT7, and 2 kb promoter sequence of *GLU6* and *ALB13* was cloned in pABAi vector (TaKaRa Bio; USA) to form bait construct. After linearising with FastDigest *BstB*I restriction enzyme (Thermoscientific^TM^; USA), linearized pABAi bait, *pGLU6* and *ALB13* was transformed in Y1H gold yeast strain using EZ-Yeast^TM^ transformation kit (MP Biomedicals; USA), screened on SD/-Ura and growth was checked on different concentration of Aureobasidin A (AbA) to determine the minimal inhibitory concentration. To confirm the binding specificity of ZOS5-09 with the *GLU6* and *ALB13* promoter, ZOS5-09_pGADT7 with *pGLU6_pABAi* were co-transformed into the Y1H gold yeast strain using EZ-Yeast^TM^ transformation kit (MP Biomedicals; USA). Empty pGADT7 and pABAi vectors were used as controls. Transformed yeast cells were cultured on SD/-Leu/-Ura selective medium containing 200 ng/ml AbA for 4 days at 30 °C. To find the region of 2 kb *GLU6* promoter responsible for binding to ZOS5-09, overlapping deletion constructs, 400 bp each with 21 bp overlap, were prepared and cloned in pABAi vector to produce *pGLU6_1*, *pGLU6_2*, *pGLU6_3*, *pGLU6_4*, *pGLU6_5*.

### Electrophoretic mobility shift assays

Promoter sequence of *GLU6* was checked in PLANTPAN3.0 (http://plantpan.itps.ncku.edu.tw/) to find the known ZF binding site (ZF_BS). For synthesizing EMSA probes, ZF_BS binding site along with flanking sequence was used in multiples of two to produce 2X ZF_BS. EMSA was performed using Light Shift Chemiluminescent EMSA Kit (Thermo Fisher Scientific; USA), according to the manufacturer’s protocol. GST_ZOS5-09 recombinant protein and *GLU6_4* probe were used for the analysis. Pierce™ Biotin 3’ End DNA Labeling Kit (Invitrogen^TM^, USA) was used for biotin labeling of the probe. Synthetic double-stranded probes were used at a concentration of 25 fmol mL^−1^, and 50-, 100-, fold levels of nonbiotinylated probe were used as the competitor. The mutated competitors were generated by replacing CACT to GTTC in the binding site. GST_ZOS5-09 was incubated with biotin-labeled probe and/or competitor probe in the binding buffer and dI.dC at room temperature for 30 mins. The protein–probe complexes were separated by 6% non-denaturing polyacrylamide gel electrophoresis at 80-100 V for 1.5 h, transferred to a nylon membrane at 85 mA for 1 h. The nylon blot was UV-cross-linked for 1200 sec using 254 nm UV light. The protein-probe complexes were detected using a Chemiluminescent Nucleic Acid Detection Module Kit (Thermo Fisher Scientific; USA), according to manufacturer’s protocol.

### In planta reporter effector assay in Nicotiana benthamiana

To study repression property, effector plasmid, MADS29_DLN construct was prepared by fusing DLNSPP coding sequence of *ZOS5-09* to the reverse primer of *MADS29*. MADS29_DLN and full length coding sequence of *MADS29* were cloned into the entry vector pENTR^TM^/D-TOPO^®^ and then recombined into pEARLYGATE201 vector under the control of *CaMV35S* promoter (Addgene, USA) by LR reaction using Gateway^TM^ LR Clonase^TM^ II Enzyme mix (Invitrogen^TM^; USA). For the reporter plasmid, 666 bp promoter fragment from *Cys-prt* was cloned into the pMDC164 vector in-frame with GUS (*β-glucuronidase*) reporter gene (Curtis and Grossniklaus, 2003). Effector and reporter plasmids were transformed into *Agrobacterium* strain, EHA105 (Xu and Li, 2008). A transient co-expression of *MADS29* or *MADS29_DLN* effector constructs with *pCys-Prt* reporter construct in equimolar ratio into *Nicotiana benthamiana* leaves through *Agrobacterium*-mediated infiltration (Norkunas *et al*., 2018), was used as test. Only *pCys-Prt* reporter construct was used as control. To study binding of ZOS5-09 to *GLU6* promoter, 2 kb *GLU6* promoter was amplified from the IR64 genomic DNA, and the fragment was cloned into pMDC164 to produce *pGLU6::GUS* through entry vector, pENTR^TM^ D-TOPO^®^ (Invitrogen^TM^, USA) using Gateway™-based cloning (Invitrogen^TM^; USA). The coding sequences of *ZOS5-09* was amplified and cloned in pEARLYGATE201. *Agrobacterium* strain EHA105 carrying the reporter plasmid (*pGLU6*::GUS) and the effector plasmids (*pCaMV35S::ZOS5-09*) was cultured and infiltrated into *N.benthamiana* leaves. Infiltrated leaves were harvested after three days of infiltration and RNA was isolated using TRIzol Reagent^®^ Solution (Invitrogen^TM^; USA). The reporter expression was checked in test and control with six biological replicates and three technical replicates. The reporter expression was normalized with *HptII* and *blpR (Basta)* expression confirmed adequate effector activity. Student’s t test was used to assess for significant differences and GraphPad Prism v7 (https://www.graphpad.com/scientific-software/prism/) was used to make the dot-whisker plot.

### Protein interactions in yeast by yeast two-hybrid assays

For yeast two-hybrid assays, full-length coding sequence of *ZOS5-09* and *ZOS5-09* with DLN motif deleted (ZOS5-09_ΔDLN), were amplified and cloned in pGADT7 and pGBKT7 vectors (Takara™ Bio USA, Inc), thus, forming ZOS5-09_AD or BD and ZOS5-09_ΔDLN_AD or BD constructs using restriction-based cloning. Yeast strain AH109 was used to carry out yeast two-hybrid assays using the Matchmaker Gold system (Takara™ Bio USA, Inc). Histone deacetylases, i.e., HDT701, HDA704, HDA706, HDA709 and SRT701, genes (Hu *et al*., 2009) were cloned in pGADT7 and pGBKT7 vectors using restriction-based cloning and *HDT702* were cloned in pGADT7-GW and pGBKT7-GW vectors (Addgene; USA) using Gateway™-based cloning (Invitrogen^TM^; USA). All primers are listed in Table S1.

EZ-Yeast^TM^ transformation kit (MP Biomedicals; USA) was used to co-transform pGADT7 or pGADT7-GW and pGBKT7 or pGBKT7-GW plasmid constructs into AH109 yeast strain and plated on synthetic drop-out (SD) medium lacking leucine and tryptophan. Positive interactors were detected by incubating transformed yeast cells grown on SD/-Leu/-Trp/-His/-Ade and SD/-Leu/-Trp/-His/-Ade with 80 mg/L X-α-gal (5-bromo-4-chloro-3-indolyl-β-D-galactopyranoside) medium. Co-transformed T-antigen_pGADT7 and p53_pGBKT7 in yeast cells were used as the positive control while co-transformed T-antigen_pGADT7 and Lam_pGBKT7 were used as the negative control.

### Protein expression and *in vitro* pull down assays

Full-length coding sequences of *ZOS5-09* were cloned in GST (glutathione-S-transferase) tagged vector, pGEX4T-1 (Addgene; USA) and *HDA704* and *OsTPL* were cloned in MBP (maltose binding protein) tagged vector, pMALc2x (Addgene; USA) by restriction based cloning to produce GST_ZOS5-09, MBP_HDA704, MBP_OsTPL protein expression constructs, respectively. Primers used in this study are listed in Table S1.

GST and MBP-tagged recombinant proteins were expressed in competent *Escherichia coli* strain BL21 and purified using glutathione agarose beads (G-biosciences^®^; USA) and maltose beads (G-biosciences^®^; USA), respectively. For pull down assay, *in vitro* expressed GST fusion proteins or GST (control) was bound to glutathione agarose beads, overnight, followed by washing with 1X PBS (0.14 M NaCl, 2.7 mM KCl, 10.1 mM Na_2_HPO_4_, and 1.8 mM KH_2_PO_4_). After the washing, MBP fused proteins or MBP (control) were added for 4 h and washing was done with wash buffer (30 mM Tris–HCl, 50 mM NaCl, pH 7.5). Bound protein complex was eluted using 2.8 mg/ml reduced glutathione (Sigma-Aldrich^®^; USA). Eluted proteins were separated on SDS-PAGE gel and analyzed by immunoblotting with anti-GST antibody (1:4000; G7781-Sigma-Aldrich^®^) and anti-MBP antibody (1:4000; M6295-Sigma-Aldrich^®^). Empty pGEX4T-1 and empty pMAL-c2x were used as controls in all reactions.

### Co-IP assay

Co-IP assay was performed using Dynabeads™ Protein A Immunoprecipitation Kit (ThermoScientific^TM^, USA). For the experiment, ZOS5-09 was cloned in pSITE-3CA vector in-frame with the N-terminal YFP tag and HDA704 was cloned in pGWB420 vector in-frame with the C-terminal MYC tag. These constructs were transformed in *Agrobacterium* strain EHA105 for infiltration in three-week old *N. benthamiana* leaves. In control, empty MYC tagged vector and YFP_ZOS5-09 were co-transfected whereas in case of test, YFP_ZOS5-09 and HDA704_MYC were co-transfected in equal molar ratios. The infiltrated plants were kept at 28°C in the dark for 48 hrs and leaf samples were then harvested. Total protein from 100 mg of leaf samples were extracted using 3-4 ml of protein extraction buffer (20 mM Tris-Cl, pH, 7.5, 150 mM NaCl, 1mM EDTA, 1mM DTT and 1mM PMSF). The cell lysate was centrifuged at 100g for 10 min at 4°C. The supernatant was separated and incubated with 50 μl of Dynabeads Protein A (pre-equilibrated with 1-3 µg of anti-MYC antibody) for 4 h at 4°C and washed three times with wash buffer and finally eluted using 20 µl elution buffer (given in kit). These samples were then analyzed by immunoblotting with anti-YFP antibody (1:2500; G7781-Sigma-Aldrich^®^) and anti-MYC antibody (1:10000; M6295-Abcam).

### Immunoblot analysis to show deacetylation

Total protein was isolated from 100 mg calli from wild type and 5-09_OE using a standard protocol (Lang *et al*., 2013). 40 µg of total protein was separated on 10% SDS-PAGE gel (Kresge *et al*., 2006). For Ponceau-S staining, blot was incubated in Ponceau-S solution (40% methanol, 15% acetic acid, 0.25% Ponceau-S). Immunoblot analysis was performed using α-Histone H4 (acetyl K5 + K8 +K12 + K16) antibody (1:10000; ab177790-Abcam).

### Statistical analysis

One-tailed student t-test was used to calculate statistical significance assuming equal variance for two samples in Microsoft Excel^®,^ p<0.05 was considered as significant. All the experiments were performed in biological replicates, as mentioned for each experiment.

### Accession numbers

The rice sequences were found from Rice Genome Annotation Project with following locus Ids: *AGPS1* (LOC_Os09g12660), *AGPL1* (LOC_Os03g52460), *ALB3* (LOC_Os03g55740.1), *ALB13* (LOC_Os07g11650), *BeIIb* (LOC_Os02g32660.1), *CDKA1* (LOC_Os03g02680), *CDC20* (LOC_Os03g02680), *CYCA* (LOC_Os12g31810), *CYCB* (LOC_Os01g59120), *EXPA1* (LOC_Os04g15840), *EXPA2* (LOC_Os01g60770), *EXPA3* (LOC_Os05g19570), *EXPA4* (LOC_Os05g39990), *FLO7* (LOC_Os10g32680), *GBSSI* (LOC_Os06g04200), *GLB4* (LOC_Os03g57960), *GLU6* (LOC_Os02g15090), *GLU19* (LOC_Os09g37976), *HDA704* (LOC_Os07g06980), *HDA706* (LOC_Os06g37420), *HDA709* (LOC_Os11g09370), *HDA711* (LOC_Os04g33480), *HDT701* (LOC_Os05g51830), *HDT702* (LOC_Os01g68104), *MCM3* (LOC_Os05g39850), *MST4* (LOC_Os03g11900), *PRO16* (LOC_Os06g31070.1), *PRO18* (LOC_Os07g10580.1), *SSI* (LOC_Os06g06560), *SSIIIa* (LOC_Os08g09230), *SRT701* (LOC_Os04g20270), *TPL* (LOC_Os08g06480), *ZOS5-09* (LOC_Os05g38600).

## Acknowledgements

The authors thank central instrumentation facilities (CIF) of NIPGR for usage of confocal, stereo and scanning electron microscopes. We thank the NBT e-library consortium for providing online access to research articles. A.M. acknowledges University Grants Commission for JRF and SRF fellowships. P.J. and F.Q. are thankful to Council of Scientific and Industrial Research, for JRF and SRF fellowships.

## Authors contributions

P.A. designed and supervised the experiments. P.J., F.Q. and A.M. performed experiments related to ZOS5-09. A.V. generated knock-down plants of ZOS5-09. A.K.T. provided scientific inputs during investigation and manuscript preparation. P.A. and P.J. analyzed the data, prepared the figures and wrote the article. All authors have read and approved the article.

## Data availability

All data supporting the findings of this study are available within the paper and within its supplementary materials published online.

## Short legends for Supporting Information

Fig. S1. Expression study of *ZOS5-09* through qRT-PCR and GFP fluorescence analysis.

Fig. S2. Localization of ZOS5-09 in *Nicotiana benthamiana*.

Fig. S3. Transactivation and repression study of ZOS5-09.

Fig. S4. Lethal phenotype of *ZOS5-09* overexpression plants.

Fig. S5. Generation of 5-09_SOE plants and their phenotyping in T_0_ generation.

Fig. S6. CRISPR Cas9-mediated mutagenesis in *ZOS5-09*.

Fig. S7. Segregation of the mutation and off-target analysis in 5-09_KO lines.

Fig. S8. Germination percentage of *ZOS5-09* knock-down T_3_ grains and knock-out T_2_ grains.

Fig. S9 Analysis of seed parameters of 5-09_KO in T_1_ grains.

Fig. S10. Analysis of reproductive parameters of 5-09_KD in T_1_ grains.

Fig. S11. Phenotyping of *ZOS5-09* knock-down (KD) and knock-out (KO) plants.

Fig. S12. Expression of *LOC_Os01g62190* in *ZOS5-09* knock-down (KD) and knock-out (KO) plants.

Fig. S13. Knock-down and knock-out of *ZOS5-09* grains show increased cell expansion in rice spikelet hull.

Fig. S14. Endosperm sections of 5-09_KD and 5-09_KO grains.

Fig. S15. *ZOS5-09* influences starch accumulation.

Fig. S16. Downstream interactions of ZOS5-09.

Fig. S17. Interaction of ZOS5-09 with OsTPL.

Fig. S18. Interactions of ZOS5-09 with histone deacetylases.

Fig. S19. Y2H assay of ZOS5-09 with HDA704.

Table S1. Primers used in the study

Table S2. Comparison of expression of ACT1, UBQ5, GAPDH reference genes

