## Supplemental Table S1 for "A zinc finger transcriptional repressor ZOS5-09 regulates rice grain size, starch and protein biosynthesis"

| **Table S1. Primers used in the study** | | | | | |
| --- | --- | --- | --- | --- | --- |
| **Primers used for promoter, localization and repression studies** | | |  |  |  |
| **Constructs** | | **Sequence (5’ to 3’)** | **Purpose** | **Amplicon size** | **Restriction enzymes** |
| pZOS5-09 | | F- CACCGTCCCCTTGTATAAAGTTG | For amplification of promoter region | 1618 bp | - |
|  |  | R-TCCACCTCGCGCAAATGTAC |  |  | - |
| YFP-5-09 | | F-CACCATGGCGAGTGGGCAACGA | Cloning in pSITE-3CA | 522 bp | - |
|  |  | R-CACGAGCAGATTAAGCAGATGG |  |  | - |
| YFP-(ΔNoRS)-5-09 | | F-CACCATGGCGAGTGGGCAACGA | Cloning in pSITE-3CA | 504 bp | - |
|  |  | R-ATGGGGCTCCTGTTCCGCACGCTC |  |  | - |
| MADS29_pEARLYGATE201 | | F-CACCATGGGGCGCGGCAAGATCGAG | For cloning in pEARLYGATE201 | 780 bp | - |
|  |  | R- CCACAGCTGCAGGCCGTGGCCGGC |  |  | - |
| MADS29+ZOS5-09_DLN_pEARLYGATE201 | | F-CACCATGGGGCGCGGCAAGATCGAG | For cloning in pEARLYGATE201 | 808 bp | - |
|  |  | R-CGGCGGGCTGTTCAAGTCCCACAGCTGCAG |  |  | - |
| pCys-Prt_pMDC164 | | F-CACCTGGTTTATTGGGTTCCGATCA | For cloning in pMDC164 vector | 407 bp | - |
|  |  | R-TTCCCTACCACATACTCACAT |  |  | - |
| **Primers used for DNA-protein interaction studies** | | |  |  |  |
| **Constructs** | **Sequence (5’ to 3’)** | | **Purpose** | **Amplicon size** | **Restriction enzymes** |
| pGLU6_pABAi | F-CGCGGAGCTCGAAAAGAAAGGGACATTTGT | | Cloning in pABAi | 2 kb | *Sac*I |
|  | R-CGCGGTCGACAGCTATTTGGAGAAGTTAAAATTTGA | |  |  | *Sal*I |
| pGLU6_1_pABAi | F-ATGCGAGCTCGAAAAGAAAGGGACATTTGTAT | | Cloning in pABAi | 400bp | *Sac*I |
|  | R-GTCGACTTCCCACCTTCTATATCTTTT | |  |  | *Sal*I |
| pGLU6_2_pABAi | F-GAGCTCAAAAGATATAGAAGGTGGGAA | | Cloning in pABAi | 421 bp | *Sac*I |
|  | R-GTCGACAAAGATAAAGCCAAGCACCAG | |  |  | *Sal*I |
| pGLU6_3_pABAi | F-GAGCTCCTGGTGCTTGGCTTTATCTTT | | Cloning in pABAi | 421 bp | *Sac*I |
|  | R-GTCGACAGAGAATTATAATGTGTTCTT | |  |  | *Sal*I |
| pGLU6_4_pABAi | F-GAGCTCAAGAACACATTATAATTCTCT | | Cloning in pABAi | 421 bp | *Sac*I |
|  | R-GTCGACTGTAACCTGTATTAATCCATT | |  |  | *Sal*I |
| pGLU6_5_pABAi | F-GAGCTCAATGGATTAATACAGGTTACA | | Cloning in pABAi | 421 bp | *Sac*I |
|  | R-CGCGGTCGACAGCTATTTGGAGAAGTTAAAATTTGA | |  |  | *Sal*I |
| pALB13_pABAi | F-CGCGAAGCTTTTCAATGTGTTGTACCGTTGT | | Cloning in pABAi for yeast-one hybrid assay | 2 kb | *Hind*III |
|  | R-ATCGGTCGACAGCTATACTTCGATGAGCTT | |  |  | *Sal*I |
| pALB13_1_pABAi | F-CGCGAAGCTTTTCAATGTGTTGTACCGTTGT | | Cloning in pABAi for yeast-one hybrid assay | 400 bp | *Hind*III |
|  | R-GTCGACCAAACATAAATAAAGCATAAA | |  |  | *Sal*I |
| pALB13_2_pABAi | F-AAGCTTTTTATGCTTTATTTATGTTTG | | Cloning in pABAi for yeast-one hybrid assay | 421 bp | *Hind*III |
|  | R-GTCGACTTTGGCCAATTTGGTGTCTTC | |  |  | *Sal*I |
| pALB13_3_pABAi | F-AAGCTTAGAAGACACCAAATTGGCCAAA | | Cloning in pABAi for yeast-one hybrid assay | 421 bp | *Hind*III |
|  | R-GTCGACGTTGATGGACAACTACCTCAC | |  |  | *Sal*I |
| pALB13_4_pABAi | F-AAGCTTGTGAGGTAGTTGTCCATCAAC | | Cloning in pABAi for yeast-one hybrid assay | 421 bp | *Hind*III |
|  | R-GTCGACTGTTAGAAAAAAAAAGTTCCG | |  |  | *Sal*I |
| pALB13_5_pABAi | F-AAGCTTCGGAACTTTTTTTTTCTAACA | | Cloning in pABAi for yeast-one hybrid assay | 421 bp | *Hind*III |
|  | R-ATCGGTCGACAGCTATACTTCGATGAGCTT | |  |  | *Sal*I |
| pGLU6_pMDC164 | F-CACCGAAAAGAAAGGGACATTTGTA | | Cloning in pMDC164 | 2 kb | - |
|  | R-AGCTATTTGGAGAAGTTAAAATTT | |  |  | - |
| **Primers used for protein-protein interaction studies** | | |  |  |  |
| **Constructs** | **Sequence (5’ to 3’)** | | **Purpose** | **Amplicon size** | **Restriction enzymes** |
| ZOS5-09_AD/BD/ pGEX4T-1 | F- GCCGAATTCATGGCGAGTGGGCAACGAG | | For cloning in pGADT7-AD, pGBKT7-BD and pGEX4T-1 vector | 522 bp | *EcoR*I |
|  | R-CGCGTCGACCACGAGCAGATTAAGCAGATGG | |  |  | *Sal*I |
| ZOS5-09_ ΔDLN_AD/BD | F- GCCGAATTCATGGCGAGTGGGCAACGAG | | For cloning in pGADT7-AD and pGBKT7-BD | 426 bp | *EcoR*I |
|  | R-CGCGTCGACTTCACCGGCCTCCTCGACCGC | |  |  | *Sal*I |
| HD704_AD/BD/pMALc2x | F-GATCGAATTCATGGCGTCTGATATGAGGTCACTCAACAGC | | For cloning in pGADT7-AD, pGBKT7-BD and pMALc2x vector | 1869 bp | *EcoR*I |
|  | R-CAGTGGATCCGGGCTCTGCTTCATCATTCAATTTTTC | |  |  | *BamH*I |
| HD706_AD/BD | F-GATCGAATTCATGGCCTCTTCCGCGCCGTCCG | | For cloning in pGADT7-AD and pGBKT7-BD vector | 1056 bp | *EcoR*I |
|  | R-CGAGGGATCCGCCAAGCTGGCTACCAAGCTCTATCAA | |  |  | *BamH*I |
| SRT701_AD/BD | F-GATCGAATTCATGTCACTTGGCTATGCCGAGAAGCTATCC | | For cloning in pGADT7-AD and pGBKT7-BD vector | 1449 bp | *EcoR*I |
|  | R-CGTAGGATCCAACTTTCTGAGTAGCAGGGTTCAAGCC | |  |  | *BamH*I |
| HDT701_AD/BD | F-GACTGAATTCATGGAGTTCTGGGGTCTTGAAGTCAAGCCT | | For cloning in pGADT7-AD and pGBKT7-BD vector | 891 bp | *EcoR*I |
|  | R-CTTAGGATCCCTTGGCGGGGTGCTTGGCCTTCGAG | |  |  | *BamH*I |
| HDT702_GW_AD/GW_BD | F-CACCATGGAGACGACCATGGGATTCTGGGG | | For cloning in pGADT7-GW and pGBKT7-GW | 645 bp | - |
|  | R-TTTGAAGGAGTCTCTTCTTCATCC | |  |  | - |
| HD709_AD/BD | F-GATCGAATTCATGGCCACCGGCGGGAACTCGCT | | For cloning in pGADT7-AD and pGBKT7-BD vector | 1368 bp | *EcoR*I |
|  | R-GCCGGGATCCGTCATTGAGCCTGATACGCTTCG | |  |  | *BamH*I |
| HD711_BD | F-GATCCCCGGGATGCTGGAGAAAGACAGGATAGCCTA | | For cloning in pGBKT7-BD vector | 1152 bp | *Sma*I |
|  | R-CTGTGTCGACACGGGCCCCGTCCTCATGATCATTG | |  |  | *Sal*I |
| OsTPL_GW_AD/GW_BD | F-CACCATGTCGTCGCTTAGCAGGGAGCTGGT | | For cloning in pGADT7-GW and pGBKT7-GW | 3401 bp | - |
|  | R- CAGACTTCTGGTTTGTTAGCTGCTGCCGGAGC | |  |  | - |
| OsTPL_pMALc2x | F-ATGCGGATCCATGTCGTCGCTTAGCAGGGAGCTGGT | | For cloning in pMALc2x vector | 3401 bp | *BamH*I |
|  | R-GCATAAGCTTGACTTCTGGTTTGTTAGCTGCTGCCGGAGC | |  |  | *Hind*III |
| HDT702_pMALc2x | F-GCATGGATCCATGGAGACGACCATGGGATTCT | | For cloning in pMALc2x vector | 645 bp | *BamH*I |
|  | R-GTACAAGCTTCACCTTTGAAGGAGTCTCTTCTTCATCCGA | |  |  | *Hind*III |
| **Primers used for raising transgenics** |  | |  |  |  |
| **Constructs** | **Sequence (5’-3’)** | | **Purpose** | **Amplicon size** | **Restriction enzymes** |
| 5-09_OE | F-GTCCGGATCCTTGACGATGGCGAGTG | | For cloning in pB4NU vector | 594 bp | *BamH*I |
|  | R-ACTTGAGCTCTCCCACTAGAATGGCATTG | |  |  | *Sac*I |
| 5-09_SOE | F-CACCATGGCGAGTGGGCAACGA | | For cloning in pG6SOE vector |  | - |
| 5-09_SOE_1 | R-CTACACGAGCAGATTAAGCAGATGGGG | | For cloning in pG6SOE vector | 525 bp | - |
| 5-09_SOE_2 | R-CTAATGGGGCTCCTGTTCCGCACGCTC | | For cloning in pG6SOE vector | 507 bp | - |
| 5-09_KD | F-CACCAGATGATCGACTTGAACAGCCCG | | For cloning in pANDA vector | 565 bp | - |
|  | R-CACTAGAATGGCATTGGAAACGCAACCG | |  |  | - |
| L5AD5 | F-CGGGTCTCAGGCAGGATGGGCAGTCT | | For amplifying tRNA seq with gRNA | - | - |
|  | GGGCAACAAAGCACCAGTGG | |  |  | - |
| L3AD5 | R-TAGGTCTCCAAACGGATGAGCGACAG | | For amplifying tRNA seq with gRNA | - | - |
|  | CAAACAAAAAAAAAAGCACCGACTCG | |  |  | - |
| SS_5-09 | F-GGAGGTTGAGCGTGCGGAACGTTTTAGAGCTAGAA | | For targeting signal sequence in ZOS5-09 | 20 bp | - |
|  | R-GTTCCGCACGCTCAACCTCCTGCACCAGCCGGGAA | |  |  | - |
| ZF_5-09 | F-GCGAGTGGGCAACGAGCGGTGTTTTAGAGCTAGAA | | For targeting zinc finger sequence in ZOS5-09 | 20 bp | - |
|  | R-ACCGCTCGTTGCCCACTCGCTGCACCAGCCGGGAA | |  |  | - |
| **Primer used for screening transgenics** | | |  |  |  |
| **Gene** | **Sequence (5’-3’)** | | **Purpose** | **Amplicon size** |  |
| *OsActI* | F-TCCATCTTGGCATCTCTCAG | | As endogenous control | 412 bp | - |
|  | R-GTACCCTCATCAGGCATCTG | |  |  | - |
| *OsHptII* | F-TCTACACAGCCATCGGTCCAG | | For amplification of *HptII* | 850 bp | - |
|  | R-GATGTAGGAGGGCGTGGATATG | |  |  | - |
| *U3P* | F- AGCCTTTCAGGACATGTATTG | | For target screening of CRISPR- targeted plants | 321 bp | - |
|  | R-CTTGACCCGAATTTGTGGAC | |  |  | - |
| *Cas9* | F-AAGCTGATCGCCAGAAAGAA | | For amplification of Cas9 | 549 bp | - |
|  | R-CTTATCCCGGTGCTTGTTGT | |  |  | - |
| *ZF_509* | F-AGTTCTCACCAAGTTCGCGTCCTGCTCC | | For cloning ZOS5-09 mutated sequence in pJET | 400 bp | - |
|  | R-ATATCCGCTGGTTGTTGGTAGCCCCCAC | |  |  | - |
| *SS_509* | F-ATAATGCGCCGTGTGCGGGGTCGAGTT | | For cloning ZOS5-09 mutated signal sequence in pJET | 400 bp | - |
|  | R-CAGCGATATCCAGATTATGTCCTATCCAA | |  |  | - |
| *LOC_Os08g01850.1* | F-ACTAATGGGGAACGCGTTCCGGTGCATG | | To check offtarget in ZF5-09 CRISPR- targeted plants | 219 bp | - |
|  | R-ATATGTCGATGCGGCGCGACATGGAGCC | |  |  | - |
| *LOC_Os06g49760.1* | F-TATACAGTACCCGGCGGTGTGCGTGCAG | | To check offtarget in ZF5-09 CRISPR- targeted plants | 155 bp | - |
|  | R-ATATAGGGCCGCATATGTGCCCCACGTA | |  |  | - |
| *LOC_Os02g34750.1* | F-TATCGAGGAGCCGCCGTTGTTGTGGAGG | | To check offtarget in SS5-09 CRISPR- targeted plants | 203 bp | - |
|  | R-TACAAGAGTGGCACCCCCTGCCGCTCTC | |  |  | - |
| *LOC_Os02g39200.1* | F-TGCAAATATATAGCTACAAGGAGG | | To check offtarget in SS5-09 CRISPR- targeted plants | 107 bp | - |
|  | R-TGCACAGCAAGATATCGTCCAAGTTCC | |  |  |  |
| **Primers used for qRT-PCR** | | |  |  |  |
| **Gene name** | **Sequence (5’-3’)** | | **Purpose** |  |  |
| *ALB3* | F-AAACTTTGGGCATGGGTAGCT | | To check expression by qRT-PCR | - | - |
|  | R-GCAGGAGAGCAATTGGTTGTG | |  | - | - |
| *ALB13* | F-GATCGATCGAGAGTTTGTCTTCAC | | To check expression by qRT-PCR | - | - |
|  | R-CTTGGGATTGGGAGTGCAA | |  | - | - |
| *PRO16* | F-CAGCAGCAGCCGTTTATGC | | To check expression by qRT-PCR | - | - |
|  | R-TGCCTCACGAACTCATTGCA | |  | - | - |
| *PRO18* | F-CAGGCTGGTAGCGCAACA | | To check expression by qRT-PCR | - | - |
|  | R-CACAATCGCCTGAACGCTACT | |  | - | - |
| *GLB4* | F-CGGGTTCTGAGTGGGAAATC | | To check expression by qRT-PCR | - | - |
|  | R-TTGTTGGAGAAGTACGGACTCTTG | |  | - | - |
| *GLU6* | F-GTTGTTGCACTTCCGGCTAGT | | To check expression by qRT-PCR | - | - |
|  | R-AGCCGGTGTATCACCACCAT | |  | - | - |
| *GLU19* | F-CCGGTACTGTACGTCCATGTGT | | To check expression by qRT-PCR | - | - |
|  | R-CGTCGCGTGCTGAAAACA | |  | - | - |
| *GBSSI* | F-TCTGCAACGACTGGCACACT | | To check expression by qRT-PCR | - | - |
|  | R-CCATTGGGCTGGTAGTTGTTC | |  | - | - |
| *MST4* | F-TTCGGCTACGACGTCGGTAT | | To check expression by qRT-PCR | - | - |
|  | R-AACTCGCGCAGGAAGTCATC | |  | - | - |
| *SSIIIa* | F-TCGGAAAACCGGAGGACTT | | To check expression by qRT-PCR | - | - |
|  | R-CGAGCCCGGTCTTTGTCAT | |  |  |  |
| *APS1* | F-TGCTTAAGCTTCTCCGTCAAAAT | | To check expression by qRT-PCR | - | - |
|  | R-CAGGAATAACCTCGCTTCCAAA | |  | - | - |
| *FLO7* | F-AATTTGCTGAACGCCTGGTACT | | To check expression by qRT-PCR | - | - |
|  | R-GTTCGTTGACCTCATGATTTGG | |  | - | - |
| *APL1* | F-CGGAGCGTACAGGCTGATC | | To check expression by qRT-PCR | - | - |
|  | R-TTGTTGATGCCGCTGTTTATG | |  | - | - |
| *BeIIb* | F-CCTGGGTGATGCGGACTATC | | To check expression by qRT-PCR | - | - |
|  | R-CATCGCGCGGTCAAACTC | |  | - | - |
| *ZOS5-09* | F-CCGCCGGAGATCGACTTGAA | | To check expression by qRT-PCR | - | - |
|  | R-CACGCTCAACCTCCTGATCACCTTCA | |  | - | - |
| *OsACT1* | F-CAGCCACACTGTCCCCATCTA | | To check expression by qRT-PCR | - | - |
|  | R-AGCAAGGTCGAGACGAAGGA | |  | - | - |
| *UBQ5* | F-CACTTCGACCGCCACTACT | | To check expression by qRT-PCR | - | - |
|  | R-CCTAAGCCTGCTGGTT | |  | - | - |
| *GAPDH* | F-AAGCCAGCATCCTATGATCAGAT | | To check expression by qRT-PCR | - | - |
|  | R-CGTAACCCAGAATACCCTTGAGT | |  | - | - |
| *NbEF1α* | F-AGCTTTACCTCCCAAGTCATC | | To check expression by qRT-PCR | - | - |
|  | R-AGAACGCCTGTCAATCTTGG | |  | - | - |
| *BlpR* | F-GCTCTACACCCACCTGCTGAA | | To check expression by qRT-PCR | - | - |
|  | R-ACAGCGACCACGCTCTTGA | |  | - | - |
| *GUS* | F-CAAAGCGGCGATTTGGAA | | To check expression by qRT-PCR | - | - |
|  | R-GCCAGGCCAGAAGTTCTTTTT | |  | - | - |
| *HptII* | F-CCGCAAGGAATCGGTCAAT | | To check expression by qRT-PCR | - | - |
|  | R-GATCAGCAATCGCGCATATG | |  |  |  |
| *CDKA1* | F-GGTTTGGACCTTCTCTCTAAAATGC | | To check expression by qRT-PCR | - | - |
|  | R-AGAGCCTGTCTAGCTGTGATCCTT | |  | - | - |
| *CYCA* | F-AGGTTGTCAAGATGGAGAGCGA | | To check expression by qRT-PCR | - | - |
|  | R-CGCTTTTTGTCTTCCTGGCA | |  | - | - |
| *CYCB* | F-CTCAAGGCTGCACAATCTGACA | | To check expression by qRT-PCR | - | - |
|  | R-GCATTGACGGCTGGAATTTG | |  | - | - |
| *CDC20* | F-TCGAATCACCTGTTTGTTGGC | | To check expression by qRT-PCR | - | - |
|  | R-TGGAGACAATCCAACGCAAAG | |  | - | - |
| *MCM3* | F-TTCATGCGTCACTAAATGCGAG | | To check expression by qRT-PCR | - | - |
|  | R-TGAATCTGGAAGCCCAATGTTC | |  | - | - |
| *EXPA1* | F-AGTGACGCTTCAGGAACGAT | | To check expression by qRT-PCR | - | - |
|  | R-TAGTCGCACATGATCCGGTA | |  | - | - |
| *EXPA2* | F-AGGTGGTGTCTTGTAACTTTTGTTGTTA | | To check expression by qRT-PCR | - | - |
|  | R-ATGGTCCCAAAAGCACAAGAGT | |  | - | - |
| *EXPA3* | F-TCCCCGTCAACTACAAGAGG | | To check expression by qRT-PCR | - | - |
|  | R-TCACGGTCACAAGCTCAAAG | |  | - | - |
| *EXPA4* | F-GGCGTTCTCTTCCTCCTCTT | | To check expression by qRT-PCR | - | - |
|  | R-ACCCTTGGCTGTACAGGTTG | |  | - | - |
