## Supplemental Table S2 for "A zinc finger transcriptional repressor ZOS5-09 regulates rice grain size, starch and protein biosynthesis"

| **Table S2. Comparison of expression of *ACT1*,  *UBQ5*, *GAPDH* reference genes** | | | | |
| --- | --- | --- | --- | --- |
| **Sample** | **Fold change(*UBQ5 vs Act1*)** | **t-test (*UBQ5* vs *Act1*)** | **Fold change (*GAPDH vs Act1*)** | **t-test (*GAPDH* vs *Act1*)** |
| WT | 1.124689693 | _ | 1.093456695 | _ |
| L1.1 | 2.159011 | 0.086581468 | 1.740211959 | 0.239588948 |
| L2.3 | 1.029624386 | 0.421248594 | 0.860312677 | 0.239588948 |
| ZF | 1.069581644 | 0.450388067 | 1.142206028 | 0.447094056 |
| SS | 1.547612172 | 0.198483052 | 1.410142677 | 0.249848805 |
| **Sample** | **Fold change(*UBQ5 vs Act1*)** | **t-test (*UBQ5* vs *Act1*)** | **Fold change (*GAPDH vs Act1*)** | **t-test (*GAPDH* vs *Act1*)** |
| WT | 1.035668139 | _ | 1.060930199 | _ |
| L1.1 | 47.82200739 | 0.166154792 | 2.066671723 | 0.101394793 |
| L2.3 | 0.779473971 | 0.14883533 | 0.669233181 | 0.121657774 |
| ZF | 2.218382479 | 0.151598443 | 0.44148269 | 0.069479819 |
| SS | 0.587651293 | 0.083396108 | 0.491405843 | 0.066390949 |
| **Sample** | **Fold change(*UBQ5 vs Act1*)** | **t-test (*UBQ5* vs *Act1*)** | **Fold change (*GAPDH vs Act1*)** | **t-test (*GAPDH* vs *Act1*)** |
| P1 (0-5 cm) | 1.067839954 | 0.371571384 | 2.490333688 | 0.002309729 |
| P2 (5-10 cm) | 0.975755914 | 0.473885617 | 1.79471199 | 0.157121195 |
| S1 | 0.80184148 | 0.236297945 | 2.431346195 | 0.058175756 |
| S2 | 0.813112038 | 0.03339771 | 1.486310426 | 0.166527116 |
| S3 | 1.113203846 | 0.427726864 | 0.79749934 | 0.228767789 |
| S4 | 0.969500702 | 0.448126166 | 0.721072734 | 0.081449753 |
| S5 | 1.295283042 | 0.070116929 | 0.881127794 | 0.1589476 |
| Flag leaf | 0.910295364 | 0.394715851 | 0.735518827 | 0.227885899 |
| Second leaf | 1.271283022 | 0.005857403 | 1.084343041 | 0.056719294 |
| Leaf sheath | 0.908586737 | 0.3327413 | 0.759830209 | 0.107925092 |
| Node | 0.907813971 | 0.068716282 | 0.762481276 | 0.027080278 |
| Internode | 1.534422129 | 0.202260726 | 0.860759104 | 0.171153854 |
| Root | 1.446692525 | 0.155947094 | 0.958378939 | 0.386660158 |
| Callus | 1.337431136 | 0.114138205 | 0.680667092 | 0.002102134 |
| Green box indicates batch of cDNA used for qRT-PCR in Figure 5 and 6. | | | |  |
| Blue box indicates batch of cDNA used for qRT-PCR in Figure 4. | | |  |  |
